# Genome-wide stability of the DNA replication program in single mammalian cells

**DOI:** 10.1101/237628

**Authors:** Saori Takahashi, Hisashi Miura, Takahiro Shibata, Koji Nagao, Katsuzumi Okumura, Masato Ogata, Chikashi Obuse, Shin-ichiro Takebayashi, Ichiro Hiratani

**Author notes:** These authors contributed equally to this work (Saori Takahashi, Hisashi Miura, Takahiro Shibata). Correspondence should be addressed to I.H. and S-i.T.

## Abstract

Here, we report the establishment of a <u>s</u>ingle-<u>c</u>ell DNA <u>repli</u>cation <u>seq</u>uencing method, scRepli-seq, which is a simple genome-wide methodology that measures copy number differences between replicated and unreplicated DNA. Using scRepli-seq, we demonstrate that replication domain organization is conserved among individual mouse embryonic stem cells (mESCs). Differentiated mESCs exhibited distinct replication profiles, which were conserved from cell to cell. Haplotype-resolved scRepli-seq revealed similar replication timing profiles of homologous autosomes, while the inactive X chromosome was clearly replicated later than its active counterpart. However, a small degree of cell-to-cell replication timing heterogeneity was present, and we discovered that developmentally regulated domains are a source of such variability, suggesting a link between cell-to-cell heterogeneity and developmental plasticity. Together, our results form a foundation for single-cell-level understanding of DNA replication regulation and provide insights into 3D genome organization.

## INTRODUCTION

In mammalian cells, DNA replication is regulated at the level of Mb-sized chromosomal units called replication timing domains, in which coordinated firing of multiple replication origins occurs^1^. Early-replicating regions generally coincide with euchromatic features, such as higher transcriptional competence and ‘active’ histone modifications, and are compartmentalized in the nuclear interior, while late-replicating regions exhibit heterochromatic features and are often located in the nuclear/nucleolar periphery^1,2,3,4^. Genome-wide DNA replication timing profiles are cell-type specific^5^ and correlate well with A (active) and B (inactive) subnuclear compartments, as revealed by Hi-C or <u>hi</u>gh-throughput chromosome conformation capture (3<u>C</u>)^6,7^. While the significance of DNA replication timing regulation remains unclear, DNA replication timing serves as an excellent forum in which to investigate the relationship between the 3-dimensional (3D) genome organization, cell identity, and cell fate changes during development^2,5^.

Our current view of DNA replication timing regulation is largely based on analysis of cell populations, and it is unclear whether our view still holds true at the single-cell level. For instance, even cells of the same type in a population use different cohorts of replication origins in every cell cycle, which results in highly heterogeneous origin usage from cell to cell^8,9,10^. Whether such origin usage heterogeneity results in variable replication timing of domains from cell to cell is an open question that cannot be addressed using existing technologies. Current methods to map replication timing domains genome-wide require at least several thousand S-phase cells fractionated by fluorescence-activated cell sorting (FACS) for effective enrichment of replicating DNA through immunoprecipitation of BrdU-substituted DNA (BrdU-IP), and obtaining such cell numbers requires ~10^6^ cells in one FACS experiment^11,12^. Such technical limitations preclude the analysis of single cells and rare cell types. Moreover, single-molecule analysis that directly visualizes DNA replication processes on extended DNA fibers is a powerful means to detect cell-to-cell variation in origin usage hidden in bulk measurements^8,9,10^. However, this method requires fluorescence in situ hybridization (FISH) to identify the extended DNA fiber that contains the chromosomal regions to be analyzed. Moreover, the average DNA fiber length that can be prepared using this type of analysis is generally ~400 kb^10^, making it difficult to uncover the structural properties of Mb-sized replication timing domains.

In the present study, we have successfully devised a novel single-cell DNA <u>repli</u>cation sequencing method, named scRepli-seq, that enables genome-wide mapping of replication domains in single mammalian cells and analyzed human TERT-RPE1 cells, as well as mouse ESCs, before and after differentiation. Our results provide a compelling set of evidence that significantly improves our single-cell-level understanding of DNA replication regulation and provides insights into 3D genome organization in the context of mESC differentiation.

## RESULTS

### A single-cell replication timing profiling method, scRepli-seq

Our routine genome-wide replication timing assay is based on immunoprecipitation of BrdU-substituted DNA (BrdU-IP) from early and late S-phase cell populations fractionated by FACS. Relative enrichment of early- and late-replicating DNA is analyzed genome-wide, either by CGH (<u>C</u>omparative <u>G</u>enomic <u>H</u>ybridization) microarrays or NGS (<u>N</u>ext-<u>G</u>eneration <u>S</u>equencing), which generates a genome-wide map of DNA replication timing^11,13^. We refrained from using the S/G1 copy number method^14^ because the signal-to-noise ratio was much lower than that of BrdU-IP when using CGH microarrays^11^. However, BrdU-IP is incompatible with single-cell analysis. Aiming to develop a single-cell replication profiling method, we decided to switch to a copy number-based method and directly tested how it compares with BrdU-IP when using NGS.

To distinguish early- and late-replicating DNA, we first used karyotypically stable human TERT-RPE1 cells^15^ and collected 100 cells at mid S-phase and G1-phase (control) via FACS and isolated genomic DNA (gDNA). The gDNA samples were subject to whole genome amplification (WGA; Sigma SeqPlex kit) followed by Illumina HiSeq NGS (Fig. 1a). Of approximately 4 million (4 M) NGS reads, ~50% were uniquely mapped to the reference genome. Mapped reads were counted throughout the genome in sliding windows of 200 kb at 40-kb intervals for both mid-S and G1 cells (Fig. 1b). G1 cells showed a nearly flat profile, suggesting that amplification bias during WGA was limited. In contrast, mid-S cells exhibited variability along the chromosomes indicative of replicated/unreplicated DNA (Fig. 1b). For normalization and comparison across multiple mid-S samples, 200-kb window read counts throughout the genome were corrected for mappability using G1 cells and further divided by the median read count (i.e., median centering), which generated a replication timing plot from 100 mid-S cells that very much resembled the Log_2_(early/late) plot from BrdU-IP experiments (Fig. 1b, Log_2_[(corrected mid-S)/median] and Fig. 1c, Pearson’s R=0.80).

**Figure 1.**
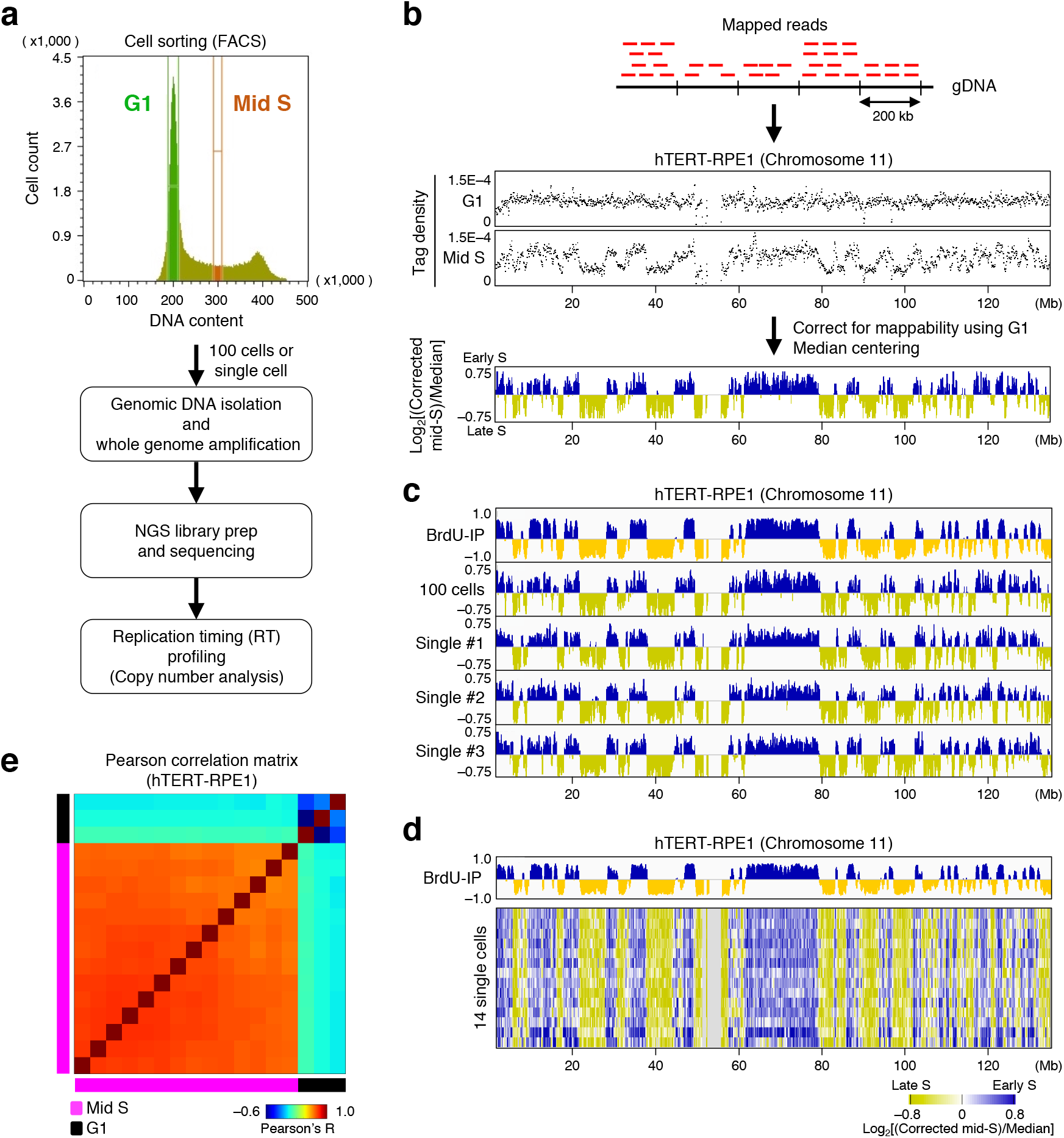
Establishment of a single-cell replication timing profiling method, scRepli-seq. (**a**) An experimental overview of scRepli-seq. A typical cell cycle profile of mammalian cells stained with propidium iodide during FACS analysis is shown, along with the mid-S and G1-phase sorting gates used. Genomic DNA samples isolated from single or 100 cells were subject to NGS followed by replication timing profiling. (**b**) Replication timing profiling by copy number analysis. Mapped NGS reads of mid-S cells were counted in sliding windows of 200 kb at 40-kb intervals to generate tag density plots (i.e., counts per window normalized by total read counts); mappability was corrected using G1 samples, and the numbers were further divided by the median read count (i.e., median centering) to generate a Log_2_[(corrected mid-S)/median] replication timing plot. Shown are human chromosome 11 data from 100 hTERT-RPE1 cells in G1 and mid S-phase. (**c**) Comparison of hTERT-RPE1 replication timing profiles derived from BrdU-IP population assay and 100 mid-S cells and three single mid-S cells using the copy number method on human chromosome 11. (**d**) The heatmap shows the replication timing of 14 single mid-S hTERT-RPE1 cells, along with BrdU-IP population data on human chromosome 11. The cells are ordered according to Pearson’s correlation coefficient values against the 100-cell data average. The gray area represents unmappable genomic regions. (**e**) Pearson correlation matrix of 14 single mid-S hTERT-RPE1 cells, along with three G1 cells.

To test whether this method is applicable to single cells, we performed the same analysis starting from single mid-S hTERT-RPE1 cells (Fig. 1a). We immediately noticed that the single-cell replication timing profiles looked similar to each other and to the profiles derived from 100 cells or the BrdU-IP method (Fig. 1c, d), which was confirmed by a Pearson correlation matrix showing a clear distinction between 14 single mid-S cells and three G1 cells (Fig. 1e). Thus, by simply combining single-cell copy number alteration (CNA) detection^16^ and fractionation of mid-S cells by FACS, we successfully established a genome-wide single-cell DNA replication profiling method, scRepli-seq.

### The megabase-sized replication domain structure is stable and conserved from cell to cell

To explore the potential of scRepli-seq, we switched to the mouse ESC system. By population BrdU-IP analysis, we first found that the replication profile of naïve mESCs grown in 2i/LIF medium [containing MEK and GSK3 inhibitors (i) and leukemia inhibitory factor (LIF)], which is a culture condition thought to maintain ‘ground-state’ pluripotency *in vitro*^17,18^, was almost indistinguishable from that of naïve mESCs grown in FBS/LIF medium^5,13^ (Supplementary Fig. 1a). Second, BrdU-IP samples analyzed by CGH microarrays and NGS were comparable (Supplementary Fig. 1a). Third, a copy number-based analysis of 200,000 mESCs without WGA produced results comparable to 100 mESCs with WGA, again negating amplification bias by WGA (Supplementary Fig. 1b). Finally, we analyzed 100 mid-S cells by the copy number method shown in Fig. 1a and successfully detected known replication timing changes^13^ identified by the BrdU-IP method upon mESC differentiation to ectoderm for 7 days (Supplementary Fig. 1c).

By analyzing 34 single mESCs using scRepli-seq, we found that despite some level of heterogeneity, the cells were similar to each other overall and clearly showed conserved Mb-sized replication domain structure (Fig. 2a). Importantly, this cell-to-cell stability of replication domains was maintained in 42 single day-7 differentiated mESCs (Fig. 2a). These results, along with the similar observation in hTERT-RPE1 cells (Fig. 1c, d), demonstrate that in mammalian cells the structure of Mb-sized replication domains and their organization is remarkably stable and conserved from cell to cell.

**Figure 2.**
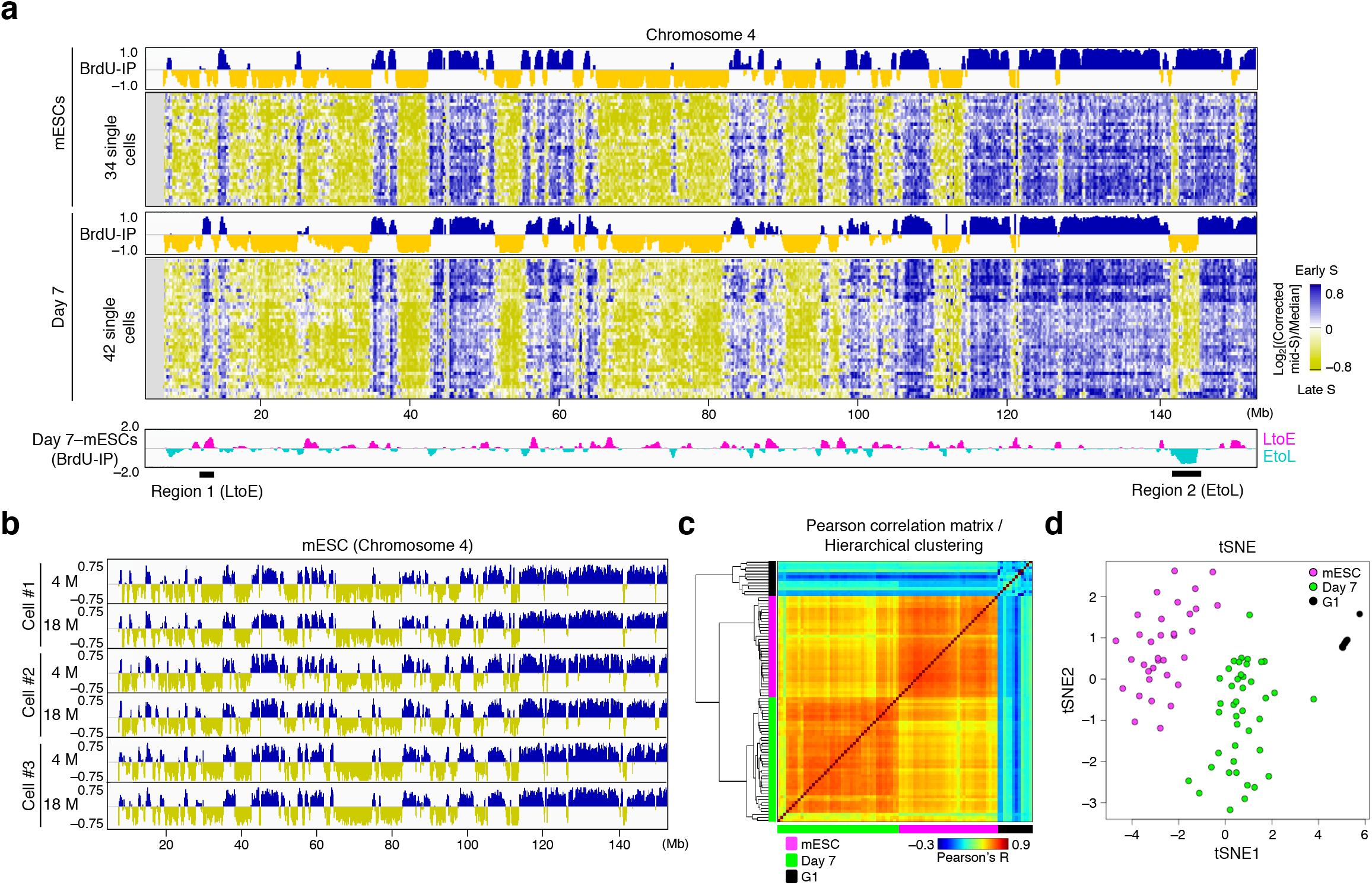
Single-cell replication profiles of mESCs before and after 7-day differentiation. (**a**) Heatmaps showing single-cell replication profiles of mESCs (34 cells) and day-7 differentiated cells (42 cells) on mouse chromosome 4, along with BrdU-IP population replication timing (RT) profiles. The plot at the bottom (in pink and light blue) shows differentials in BrdU-IP population RT data before and after differentiation (day-7 mESCs). Regions 1 and 2 are representative LtoE (late-to-early) and EtoL (early-to-late) switching regions, respectively. We found 3 out of 42 day-7 cells that did not show late replication of the *Rex2* locus (region 2), which is likely due to incomplete differentiation because two of them also exhibited early replication of both X chromosomes (the remaining one could not be binarized; see also Table 1). See also Supplementary Fig. 6. (**b**) Mouse chromosome 4 replication profiles of three single cells sequenced at low (4 M) and high depth (18 M). (**c**) Pearson correlation matrix heatmap and a hierarchical clustering tree of mid-S and G1 single-cell replication timing profiles showing distinct clusters of mESCs, day-7 differentiated cells, and G1 cells using autosomal data. (**d**) tSNE analysis showing the distinct distribution of mESCs, day-7 differentiated cells, and G1 cells using autosomal data.

### Developmental changes in replication domains can be detected at the single-cell level

While replication domain organization was stable between cells of the same type, differentiation-induced replication timing changes observed in the population data were also evident in the majority of the day-7 single cells analyzed (Fig. 2a), for both late-to-early (LtoE; e.g., Region 1 in Fig. 2a) and early-to-late changes (EtoL; e.g., Region 2 in Fig. 2a). Thus, developmental changes in replication timing can be detected in single cells. Although our temporal resolution is somewhat limited due to analysis of a single time point at mid-S, the spatial resolution reached near optimum level at only ~4 M NGS reads per cell at a 200-kb window size because an increase from 4 to 18 M reads did not improve the results in 3 independent single mESCs (Fig. 2b). Taken together, scRepli-seq can identify Mb-sized replication domains and known replication timing changes during mESC differentiation in ways comparable to population data, which is possible with only ~4 M reads per cell, making it an affordable and powerful single-cell epigenome profiling method.

Single-cell DNA replication timing profiles appeared similar among cells, but a small degree of cell-to-cell heterogeneity was also evident (Fig. 2a). However, upon hierarchical clustering of single-cell data sets derived from mESCs and day-7 cells, single mESCs formed their own cluster and could be distinguished from day-7 or G1 cells (Fig. 2c). By t-distributed stochastic neighbor embedding (tSNE), which is a dimensionality reduction technique to visualize relationships between high-dimensional data sets, we found that single mESCs and day-7 cells were again distinct from each other (Fig. 2d). Taken together, we conclude that cell-to-cell heterogeneity is confined such that single-cell replication timing profiles of a given cell type are specific and can be distinguished from those of another cell type.

### Replication timing heterogeneity is confined to a narrow S-phase time window surrounding the time of replication

Because DNA replication is a process of genome duplication, we can justify data binarization and assign ‘replicated/unreplicated’ calls to each genomic bin, which allowed us to compare the replication state of each bin across all cells, quantify cell-to-cell heterogeneity, and explore the source of the small degree of cell-to-cell heterogeneity. We used AneuFinder package in R for binarization (see Methods), which enabled us to calculate the percentage of the genome replicated in each cell (% replication score), i.e., the time each cell spent in S-phase. For instance, a ‘25% replicated’ cell is expected to have spent 25% of the entire S-phase duration at the time of fixation. As expected, many of our mid-S cells isolated by FACS (Fig. 1a) were 50 to 60% replicated, although some cells were outside this range (Fig. 3a). We excluded cells with replication scores below 40% or above 70% and focused on 33 mESCs and 35 day-7 cells, which were ordered according to their % replication scores (Fig. 3a). Binarized single-cell data sets exhibited globally similar replication profiles by visual inspection but also showed some heterogeneity (Fig. 3a; variability scores ranged from 0 to 1, where 1 indicated highest variability). Variable regions often corresponded to sequences replicated in the middle of S-phase (Fig. 3a, high variability scores correspond well with sequences replicated in mid S, i.e., 40–60% S-phase in a BrdU-IP population assay). By plotting the cell-to-cell variability scores of all 80-kb bins across the genome against their population average replication timing in % S-phase values (see Methods), we confirmed that the highest cell-to-cell variability is observed for genomic sequences replicated at approximately 60% S-phase, just after mid S-phase, in mESCs and day-7 cells (Fig. 3b, peak). Importantly, this variability peak coincided with the mean % replication score of individual cells (Fig. 3b, dotted line), indicating that sequences being replicated at the time of cell fixation showed the largest degree of cell-to-cell heterogeneity in replication state. Importantly, upon subdivision of our single-cell data sets into 3 groups with a score range of 40–50%, 50–60%, and 60–70% replication, genomic regions with the highest heterogeneity within each group corresponded to sequences replicated at 40–50%, 50–60%, and 60–70% S-phase time windows (Fig. 3c). These observations suggest that the replication timing of a specific chromosomal domain is conserved from cell to cell, although it might show slight temporal fluctuation between cells around the time of replication.

**Figure 3.**
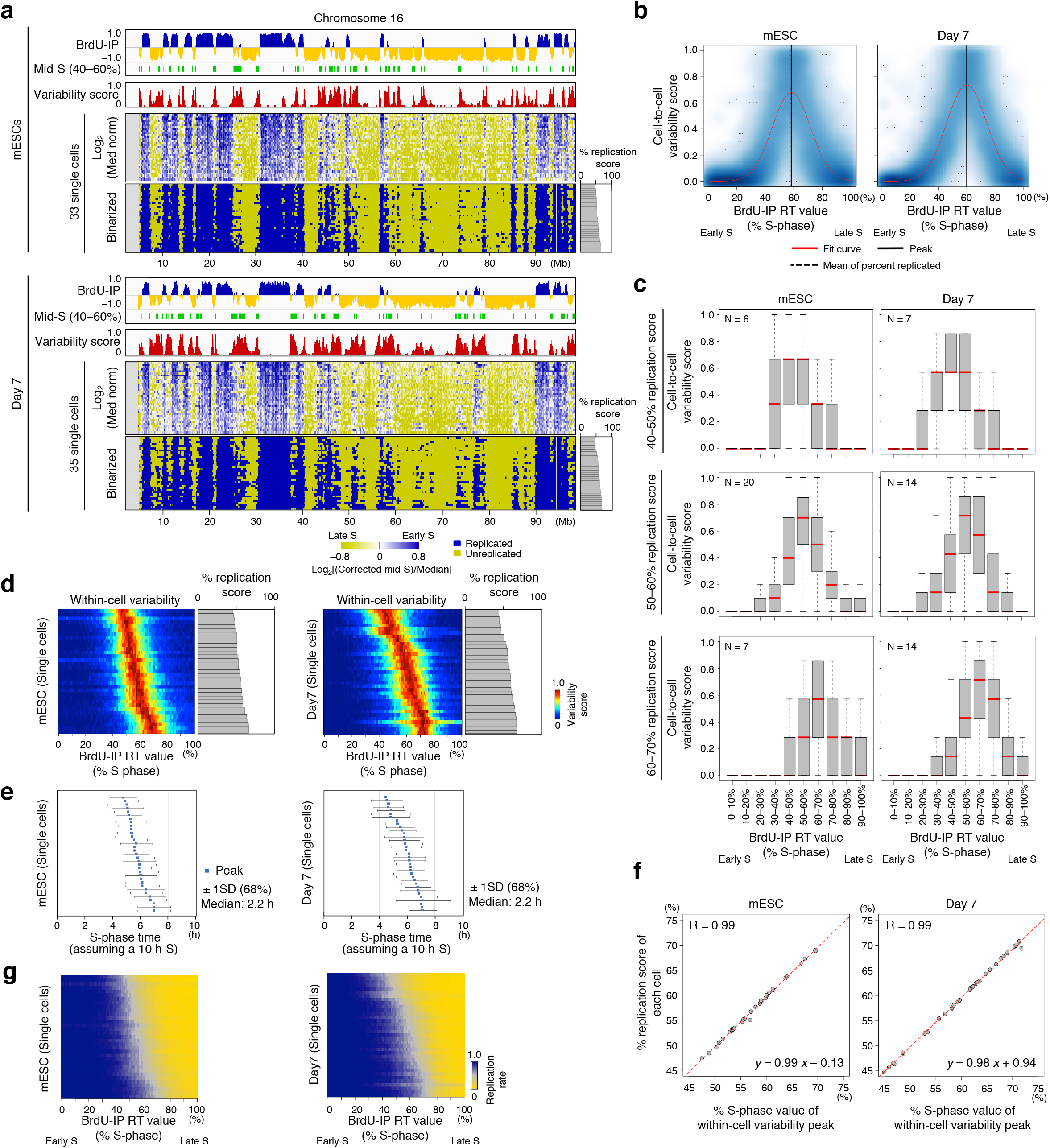
Stability and heterogeneity of replication timing regulation in single cells. (**a**) Single-cell replication profiles of mESCs and day-7 cells before and after binarization on an autosome (chromosome 4), along with BrdU-IP population RT profiles, mid-S replicating regions, and cell-to-cell variability scores. Cells are ordered according to % replication score. See also Supplementary Fig. 6. (**b**) Cell-to-cell RT variability of all 80-kb bins across the genome are plotted against their BrdU-IP RT values. Darker blue represents higher dot density. Red lines, fitted Gaussian curves; black lines, variability peaks; dotted lines, mean % replication scores of individual cells. (**c**) Relationship between cell-to-cell variability and BrdU-IP RT values. Cells were subdivided into three groups with score ranges of 40–50%, 50–60%, and 60–70% replication, and all 80-kb bins within each group were further subdivided into 10 groups according to BrdU-IP RT values. Box plots show the cell-to-cell variability distributions of each group. (**d**) We subdivided the genome into one-percentile groups of bins with similar replication timing (% S-phase scores) and measured the variability of replication state within each group in each cell in a heatmap format. Each horizontal line represents a cell. (**e**) The within-cell variability peak (blue dots) and the range of SD in each cell is shown. Percent S-phase values were converted to hours, assuming a 10-h S-phase. (**f**) Relationship between % S-phase value of the within-cell variability peak and the % replication score of each cell. (**g**) Heatmaps with a format similar to that shown in Fig. 3d showing the rate of replication within each one-percentile group of bins with similar BrdU-IP population RT values.

We next subdivided the genome into one-percentile groups of genomic bins with similar replication timing and measured the variability of replication state within each group, i.e., within-cell variability (Fig. 3d). Groups of genomic bins with high variability were confined to a narrow time window of S-phase (red bins in Fig. 3d), with 68% of the variability [± 1 standard deviation (SD) from the mean] confined to within 22% of the entire S-phase, or 2.2 h, assuming a 10-h S-phase (Fig. 3e). In addition, the % replication score of each cell (gray bar plots in Fig. 3d) positively correlated with population average replication timing values (% S-phase values in Fig. 3d) of high variability bins. Indeed, the bins with the highest variability score corresponded to bins that were just being replicated in a cell, as evidenced by the almost perfect match between the % replication score of each cell and the population average replication timing value of within-cell variability peak (Fig. 3f). Genomic bins with replication timing values earlier and later than the within-cell variability peak were mostly replicated and unreplicated in our mid-S single-cell data sets, respectively (Fig. 3g). In brief, the early-replicating region in a cell population is rarely replicated in late S-phase, even in single cells, and vice versa. We conclude that the levels of both cell-to-cell and within-cell heterogeneity in replication timing are limited, and as a result, the DNA replication timing program is highly conserved.

### Sequences subject to developmental regulation of replication timing exhibit higher cell-to-cell heterogeneity than constitutive regions in mESCs

We next thought to investigate whether sequences subject to developmental regulation of replication timing are a source of heterogeneity and show higher cell-to-cell variability than constitutively early- or late-replicating sequences. We first defined constitutively early (CE), constitutively late (CL), and developmentally regulated (D) classes of genomic bins by comparing the replication timing of 28 cell types in a manner similar to that described by Dileep et al. (Supplementary Fig. 2a)^19^. The CE- and CL-class sequences were within the first 30% (0–30% S-phase) and the last 50% (50–100% S-phase) of the genome replicated, respectively, based on a BrdU-IP population assay, while D-class sequences were replicated throughout the S-phase (Supplementary Fig. 2b). We calculated the cell-to-cell replication timing variability scores of all genomic bins within CE, CL, and D classes, subdivided them into one-percentile groups of bins with similar replication timing, and plotted the average variability score of each group (Fig. 4). In mESCs, comparison of CE versus D classes clearly indicated that the D class showed more cell-to-cell variability (Fig. 4a, c, p=0.004). Likewise, the D class showed more variability than the CL class (Fig. 4a, c, p=0.0271). Thus, developmentally regulated regions show relatively high cell-to-cell replication timing heterogeneity in mESCs, suggesting a possibility that they may be less well defined structurally and their inherent instability may confer competence for developmental regulation.

**Figure 4.**
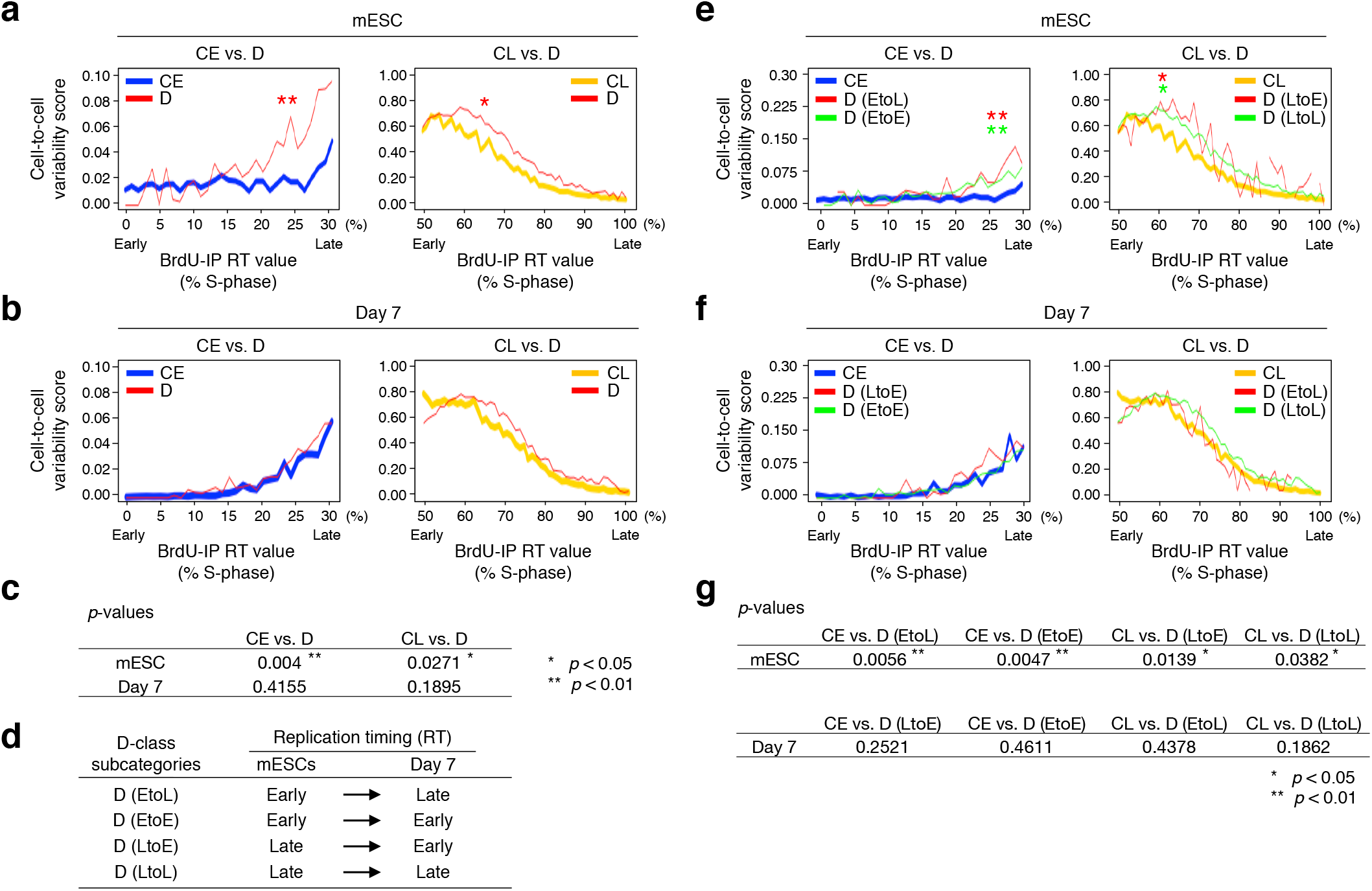
Developmentally regulated sequences exhibit higher cell-to-cell heterogeneity in RT than constitutive regions in mESCs. (**a, b**) Sequences subject to developmental regulation of replication timing (D) were compared to constitutively early (CE) or late (CL) replicating sequences for their cell-to-cell heterogeneity in mESCs (**a**) and day-7 cells (**b**). For CE-, CL-, and D-class definitions, see Supplementary Fig. 2. For each class, cell-to-cell variability scores of all 80-kb bins included in each one-percentile RT group (defined in Fig. 3d) were calculated, and the mean value was plotted. For CE vs. D and CL vs. D comparisons, sequences with BrdU-IP RT values within 0–30% and 50–100% S-phase ranges, respectively, were analyzed. Asterisks indicate statistical significance based on a permutation test (* p<0.05, ** p<0.01). (**c**) A summary of the permutation test in Fig. 4a, b. (**d**) Definitions of D-class subcategories used in Fig. 4e–g. Among the D-class bins that are early-replicating in mESCs, D (EtoL) is a subset that undergoes EtoL RT changes while D (EtoE) is a subset excluding D (EtoL). Likewise, among the D-class bins that are late replicating in mESCs, D (LtoE) is a subset that undergoes LtoE RT changes while D (LtoL) is a subset excluding D (LtoE). (e, f) Cell-to-cell heterogeneity comparison between subcategories of D and CE or CL classes in mESCs (**e**) and day-7 cells (**f**), as in Fig. 4a, b. (**g**) A summary of the permutation test in Fig. 4e, f.

Interestingly, cell-to-cell replication timing heterogeneity disappeared upon 7-day mESC differentiation (Fig. 4b, c). We further subdivided the D class into four subcategories, EtoL, LtoE, EtoE, and LtoL, based on replication timing behaviors and analyzed how their cell-to-cell variability changed before and after differentiation (Fig. 4d). The results showed that all four subcategories no longer showed higher heterogeneity on day 7, regardless of replication timing changes (Fig. 4e–g), suggesting that the observed disappearance of cell-to-cell heterogeneity during differentiation is not related to when replication timing changes occur.

### Emergence of a single late-replicating X chromosome upon mESC differentiation demonstrates the feasibility of haplotype-resolved scRepli-seq

The mESC line we analyzed was CBMS1, which is an F1 hybrid derived from a cross between CBA female and MsM/M male mice^20^. Because CBA (Mus *musculus domesticus)* and MsM/M *(Mus musculus molossinus*; henceforth MsM) are distantly related mouse subspecies, frequent SNPs/indels between these strains allowed us to generate haplotype-resolved (allele-specific) scRepli-seq data (Fig. 5a). We first generated reference genome sequences for CBA and MsM strains based on their SNP/indel information^21^ and remapped the scRepli-seq data. Although the mapping rate was not very high (~20%), likely due to our stringent criteria to define strain-specific SNPs/indels (see Methods), we nonetheless obtained haplotype-resolved data with sufficiently high resolution for further analyses (Fig. 5b, c). There are ample MsM SNPs with ~1 substitution every 100 bp^22^, and the situation should only improve with future MsM genome assembly releases (http://molossinus.lab.nig.ac.jp/msmdb/index.isp).

**Figure 5.**
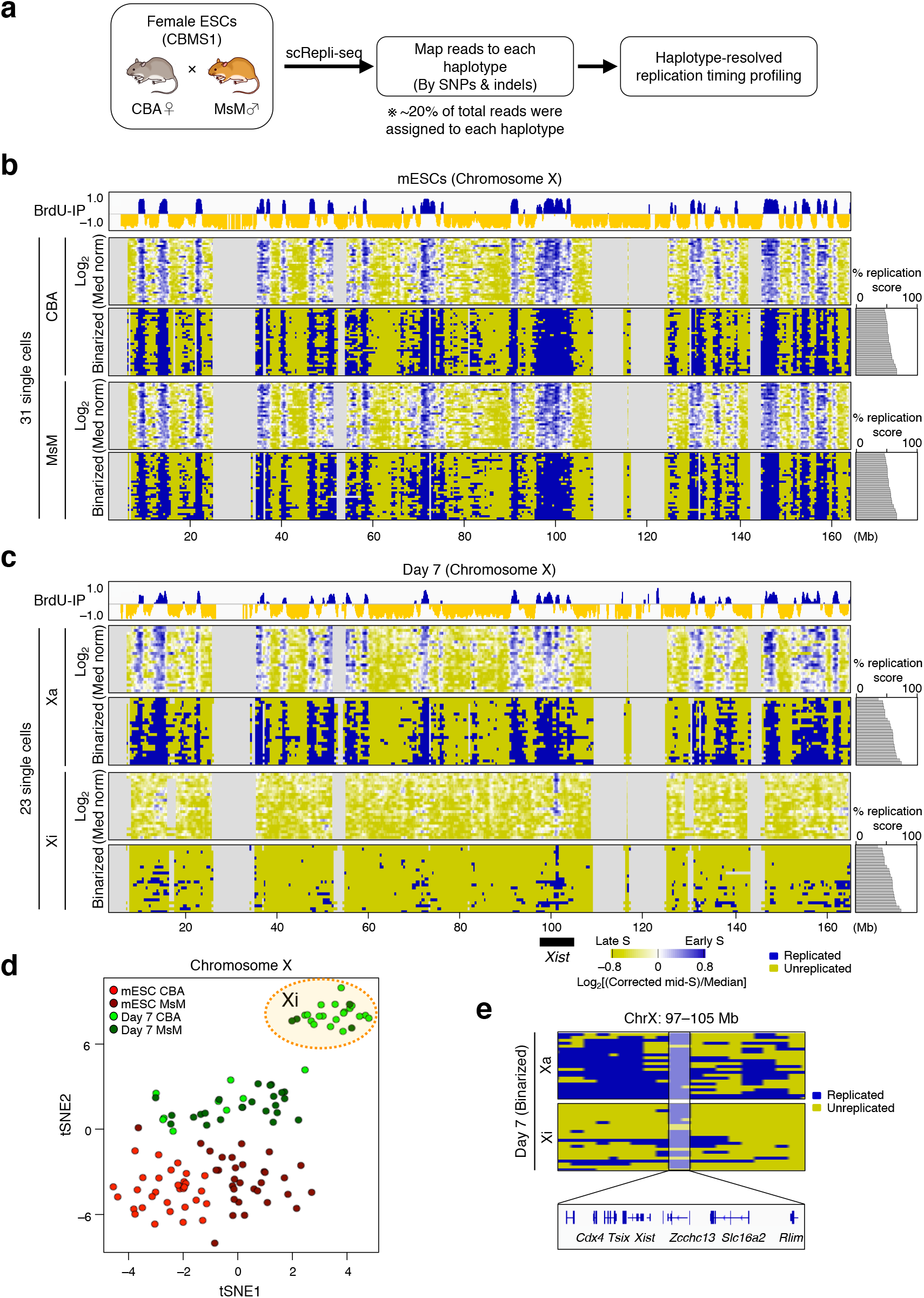
Haplotype-resolved scRepli-seq analysis of X chromosomes during mESC differentiation. (**a**) An overview of haplotype-resolved scRepli-seq analysis using CBMS1 mESCs. (**b, c**) Haplotype-resolved replication timing profiles of X chromosomes before and after binarization in mESCs (**b**) and day-7 differentiated cells (**c**). The BrdU-IP population RT and % replication of each cell are shown as well. For day-7 cells (**c**), only cells with one late-replicating X (Xi) are shown, and the other earlier-replicating counterpart was defined as the Xa. See also Supplementary Fig. 6 and Table. 1. (**d**) tSNE analysis of X chromosomes in mESCs and day-7 cells. The upper right population (highlighted in orange) corresponds to the later-replicating X in Fig. 5c. See also Supplementary Fig. 6. (**e**) A magnified view of a region surrounding the *Xist* locus in day-7 cells, which is the earliest replicating sequence on the Xi.

In female mammals, one of the two X chromosomes replicates late in somatic cells, which corresponds to the inactive X chromosome (Xi)^23,24,25^. We reasoned that the replication timing of the X chromosomes in day-7 differentiated cells would tell us whether haplotype-resolved analysis is feasible. To this end, we analyzed haplotype-resolved data derived from mESCs and day-7 cells before and after binarization, as in Fig. 3a. In mESCs, the two X chromosomes derived from CBA and MsM exhibited replication profiles very similar to each other, with both early and late-replicating domains throughout the chromosome (Fig. 5b). However, in day-7 cells, the majority had one late-replicating Xi (Fig. 5c), which is also evident in the tSNE plot showing CBA- and MsM-derived X chromosomes in mESCs and day-7 cells (Fig. 5d). Interestingly, the earliest replicating sequence on the Xi contained the *Xist* locus (Fig. 5e), a long-noncoding RNA expressed from the Xi that regulates X-chromosome inactivation (XCI), consistent with the relationship between early replication and transcription^4,26^. These results make scRepli-seq the first genome-wide methodology that can detect Xi formation at the single-cell level and also serve as a proof-of-principle for the feasibility of haplotype-resolved analysis.

### Homologous autosomes show globally similar replication timing

Fig. 6a shows the single-cell replication timing profiles of CBA and MsM haplotypes in mESCs before and after binarization. Single-cell CBA and MsM data sets resembled the population CBA and MsM data sets, respectively, and homologous chromosomes in single cells exhibited replication profiles similar to each other (Fig. 6a, Supplementary Fig. 3), which was also the case in day-7 differentiated cells (Fig. 6b, Supplementary Fig. 3). Approximately 80% of the genome exhibited the same replication state between homologous chromosomes in single cells, either both replicated (E/E) or unreplicated (L/L) (Fig. 6c). Moreover, the remaining ~20% showed one replicated and the other unreplicated (E/L), which was a pattern most frequently seen in regions that were just being replicated (Fig. 6c, red bins generally coincided with % replication scores). These bins with high E/L frequency were confined to a relatively narrow time window of S-phase (red and yellow bins in Fig. 6c), with 68% of the variability (± 1 SD from the median) confined to within 36% of the entire S-phase. In addition, the population average replication timing value of the E/L frequency peak showed a tight correlation with the % replication score of each cell (Fig. 6c). Taken together, asynchronous E/L replication patterns exist in cells, but they are often observed in regions that are just being replicated, and the unreplicated homolog will soon complete its replication as well.

**Figure 6.**
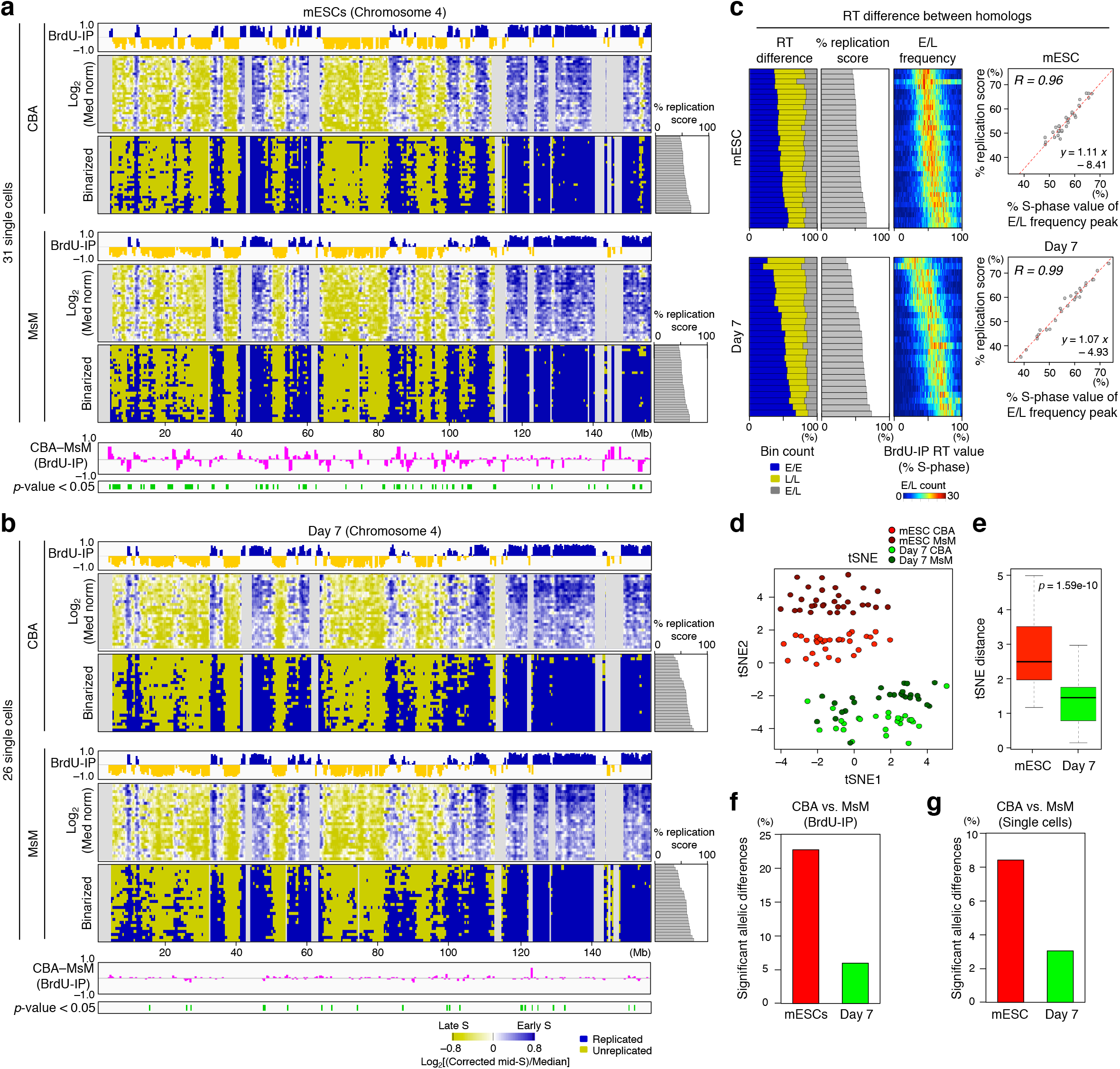
Haplotype-resolved scRepli-seq analysis of autosomes. (**a, b**) Haplotype-resolved replication timing profiles of an autosome (chromosome 4) before and after binarization in mESCs (**a**) and day-7 differentiated cells (**b**). The CBA-MsM plot presents BrdU-IP population RT differentials between the two homologs (>0, CBA earlier; <0, MsM earlier). The bar plot in green represents regions with significant allelic differences in BrdU-IP RT values based on one-way ANOVA (p<0.05). See also Supplementary Fig. 6. (**c**) Replication state of homologous regions in single cells. The leftmost bar graph shows the ratio of E/E (both replicated), L/L (both unreplicated), and E/L (one replicated) in each cell. Cells are ordered according to their %replication scores (second left). The heatmap (third from left) shows the distribution of mean E/L frequency within each of the one-percentile groups of 400-kb bins with similar BrdU-IP RT values. The rightmost scatter plot shows the relationship between % S-phase value of the E/L frequency peak (peak was defined as the mean % S-phase value of E/L frequency distribution) and % replication score. (**d**) tSNE analysis of CBA and MsM haplotypes in mESCs and day-7 cells. (**e**) Analysis of tSNE distance revealed larger within-cell allelic RT difference in mESCs than in day-7 cells. (f, g) A higher number of significant allelic differences are observed in mESCs than in day-7 cells for BrdU-IP RT data based on one-way ANOVA (p<0.05) (**f**) and averaged single-cell RT data based on Fisher’s exact test (p<0.05) (**g**).

Homologous chromosomes within a nucleus could be in different microenvironments, which is consistent with the observation that their radial subnuclear positions are frequently different^27^. Together with our observation that homologous chromosomes replicate at similar times in a given cell, it could be argued that the underlying primary sequence may be an important determinant of the replication timing. Clearly, radial positioning of chromosomes is not what determines replication timing, consistent with the inability of an acidic peptide that induces subnuclear repositioning of the targeted locus to cause a replication timing change^28^.

### Allelic replication timing differences in mESCs tend to disappear upon differentiation

By visual inspection, local allelic differences observed in mESCs using the BrdU-IP population replication timing assay were nearly absent in day-7 cells (Fig. 6a, b, compare the CBA-MsM differential plots). Consistently, tSNE analysis revealed that single-cell autosomal replication timing profiles of CBA and MsM were closer to each other in day-7 cells than in mESCs (Fig. 6d, e). We also confirmed a higher amount of significant allelic differences in mESCs than in day-7 cells in the BrdU-IP population replication timing assay (Fig. 6a, b, f; p<0.05). Furthermore, a similar trend was also observed in single-cell data sets; analysis of single-cell replication timing profiles revealed that significant allelic differences decreased substantially upon mESC differentiation (Fig. 6g). This result was reminiscent of the replication timing heterogeneity of developmentally regulated sequences in mESCs becoming less pronounced upon differentiation (Fig. 4). Taken together, between-cell and between-homolog replication timing differences in mESCs and their fate upon differentiation collectively suggest an interesting possibility that early embryonic development is accompanied by a global loss of heterogeneity in DNA replication timing.

We also analyzed whether known genomic imprinting regions showed allelic differences in replication timing. We found only one such region out of 23 genomic imprinting regions (MouseBook; www.mousebook.org/imprinting-region-list), a region on chromosome 11 containing *Grb10,* which is paternally imprinted, i.e., maternally expressed^29^. Here, the maternal homolog (CBA) is replicated earlier than the paternal one (MsM) both at the population and single-cell level (Supplementary Fig. 4). Consistent with the association between early replication and transcription^4,26^, two maternal genes are expressed as opposed to one paternal gene in this region (Supplementary Fig. 4). This asynchrony is maintained after differentiation (Supplementary Fig. 4). The fact that only one out of 23 imprinting regions showed allelic difference might seem to contradict the general view that asynchronous replication timing is associated with genomic imprinting^30,31,32,33^. However, it should be noted that the *Grb10* locus replicates at mid S-phase, which corresponds exactly to the FACS gate we used (Fig. 1a). Therefore, by sorting cells at different S-phase time windows, we think it is possible to detect replication timing asynchrony of other imprinting regions. In fact, a relatively small degree of replication asynchrony is observed for imprinting regions^34^.

### scRepli-seq and its versatility: concurrent CNA analysis and Hi-C A/B compartment predictions

Because scRepli-seq simply reads the genome, it can detect CNAs, which are often found in cancers, but we also found some CNAs during CBMS1 mESC differentiation. Female mESCs are notorious for being karyotypically unstable and losing one X chromosome to become XO^35,36,37,38^. However, we found that 31/34 (91%) CBMS1 mESCs retained two X chromosomes (XX) even after several passages, suggesting that X chromosomes are relatively stable in 2i/LIF conditions (Table 1, Supplementary Fig. 5). The remaining three CBMS1 mESCs all lost the MsM-derived X. In contrast, day-7 cells showed a much more frequent loss of one X, despite being derived from differentiating XX mESCs (Table 1, Supplementary Fig. 5). Interestingly, 13/14 (93%) XO cells had lost the CBA-derived X on day 7 (see Table 1 for more details). Among the 28 XX cells on day 7, five replicated both X chromosomes early, but three of these cells also exhibited early replication of the Rex2 locus (which normally shows an EtoL switch upon differentiation; see Fig. 2a, region 2), indicative of failed mESC differentiation. The remaining 23 cells had a late-replicating Xi, with 19 having a CBA-derived Xi and four having an MsM-derived Xi (Table 1, Supplementary Fig. 5). This result indicates that XCI was skewed (19/23=83%) in CBMS1, likely due to the *X-chromosome controlling element (Xce)* locus subtype combination effect^39^. The frequent loss of CBA-derived X only in day-7 cells (p=0.0059, Fisher’s exact test) suggests the possibility that loss of X occurs just as when it is undergoing XCI.

**Table 1.**
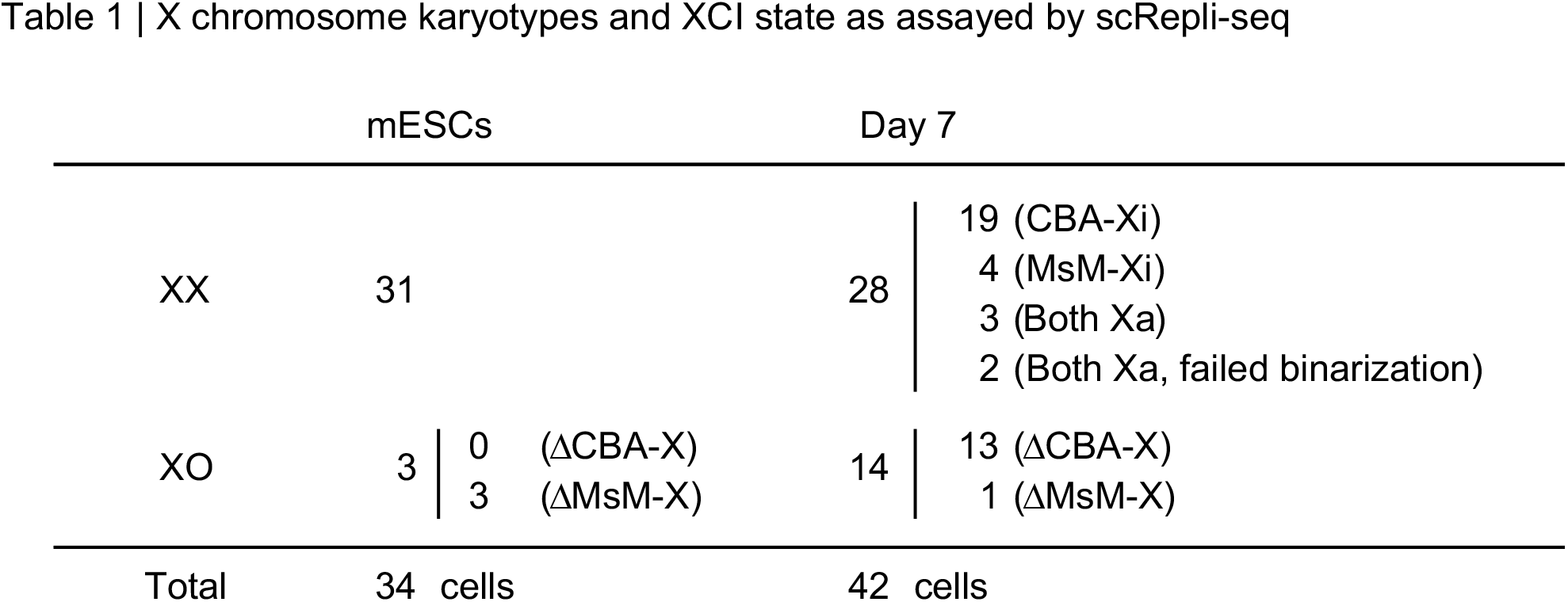
X-chromosome karyotypes and XCI states as assayed by scRepli-seq. A summary of X-chromosome karyotypes and XCI states of single cells analyzed in this study. See also Supplementary Figs. 5 and 6. Note that we included the day-7 XX^del^ cells (with a distal deletion of CBA-derived X; see also Supplementary Figs. 5) under the XO category, ΔCBA-X, for simplicity. ‘Both Xa, failed binarization’ represents samples that exhibited an Xa/Xa state based on tSNE analysis but showed binarization failures.

In addition, replication timing profiles are closely correlated with the A/B subnuclear compartments as assayed by Hi-C^7^, which was indeed the case in CBMS1 mESCs. Replication timing data sets derived from a BrdU-IP population assay, scRepli-seq, and averaged scRepli-seq before and after haplotype resolution all showed high correlations with mESC Hi-C A/B compartments (Fig. 7a, b, c), indicating that scRepli-seq allows the prediction of A/B compartments at the single-cell level. This outcome is valuable, given that although single-cell Hi-C is now feasible, low resolution still precludes identification of A/B compartments and topologically associating domains (TADs) purely from raw single-cell Hi-C data^40,41,42^. Moreover, conservation of genome-wide replication timing profiles between individual cells suggests an intriguing possibility that A/B compartment organization may also be conserved from cell to cell.

**Figure 7.**
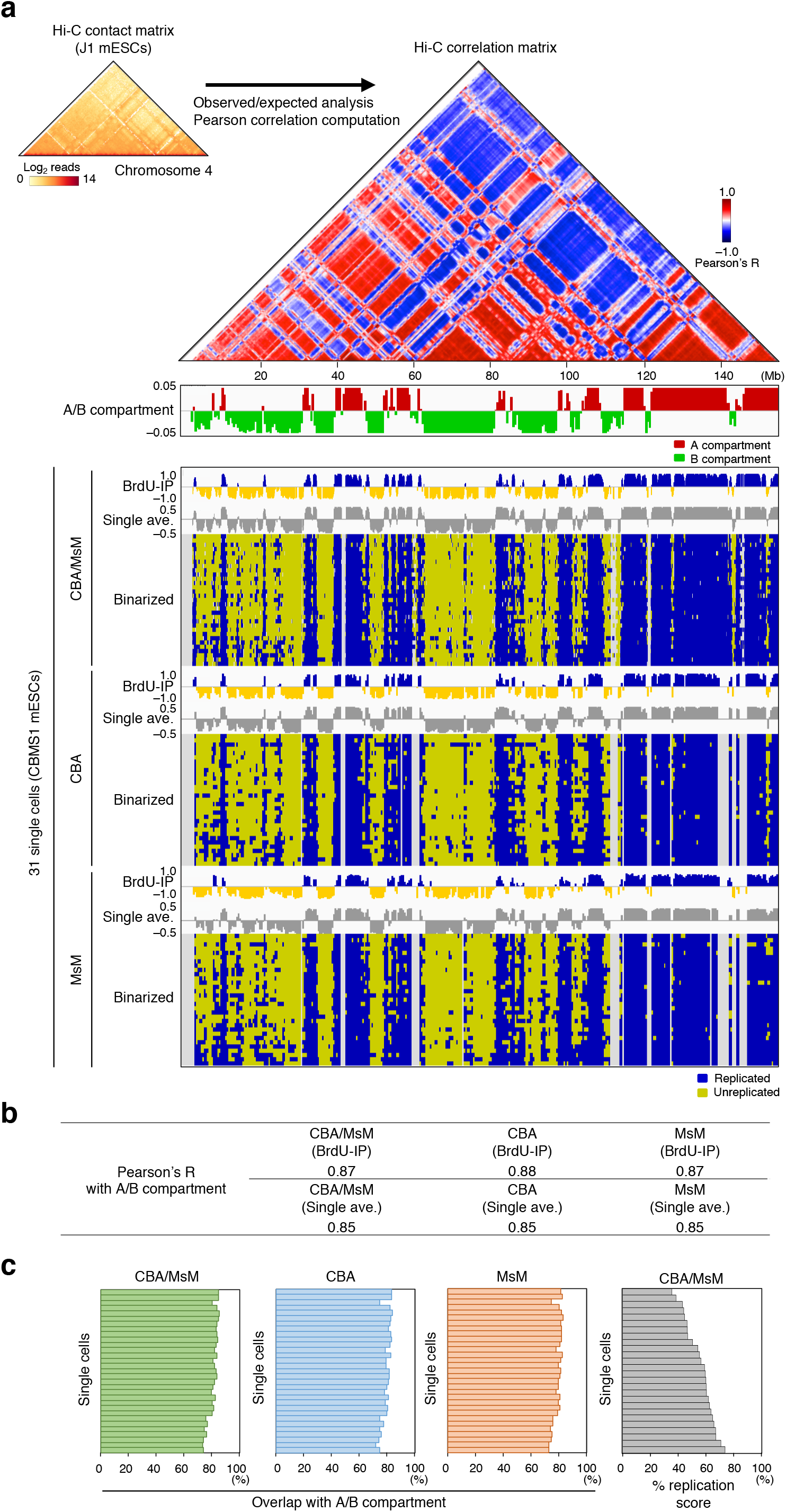
Single-cell replication timing data correlate well with Hi-C A/B compartments. (**a**) Schematics at the top show the calculation of A/B compartments from a Hi-C contact heatmap^47^ via observed/expected and Pearson correlation heatmaps. Mouse chromosome 4 is shown. Heatmaps in blue and yellow show binarized single-cell replication profiles of mESCs at 400-kb resolution on mouse chromosome 4 before and after haplotype resolution. BrdU-IP population RT profiles and averaged single-cell profiles are also shown, with and without haplotype resolution. (**b**) Pearson *R* values obtained by genome-wide comparison of A/B compartments and various RT data shown in Fig. 7a (sex chromosomes were excluded from analysis). (**c**) The rate of overlap between A/B compartments and replicated/unreplicated states in the 31 single cells shown in Fig. 7a is high and uniform from cell to cell.

## DISCUSSION

In this study, we established a genome-wide single-cell replication timing sequencing method, scRepli-seq, and analyzed human and mouse cultured cells. Our scRepli-seq results demonstrate that (1) DNA replication domains in mice and humans are stable Mb-sized structures that are highly conserved between cells and homologous chromosomes genome-wide, with only a small degree of heterogeneity; (2) DNA replication domain organization is cell-type specific even at the single-cell level; (3) developmentally regulated replication timing domains exhibit higher cell-to-cell heterogeneity than constitutive regions in mESCs, which is diminished upon differentiation; and (4) single-cell replication timing profiles are highly correlated with A/B compartments as assayed by Hi-C, suggesting an intriguing possibility that A/B compartment organization is conserved from cell to cell.

In the prevailing view, DNA replication in mammals is regulated at the level of Mb-sized domains by clusters of replication origins that fire stochastically yet nearly synchronously in each cell^2,43^. Our results are consistent with this view, but we further provide compelling evidence that domain replication timing is highly conserved from cell to cell and between homologs, which results in a conserved temporal order of genome duplication between cells (Fig. 3). This result contrasts with budding and fission yeasts, in which origin firing is stochastic and results in heterogeneous replication timing patterns from cell to cell^44,45^. To reconcile these differences, we need to assume an additional regulatory layer in mammalian cells that specifies replication timing of Mb-sized domains in a nearly deterministic manner despite stochastic origin firing, as has been suggested previously^2,46^.

What could be the identity of this additional regulatory layer? We and others have shown that DNA replication timing correlates well with A/B compartments as assayed by Hi-C^7,47^ (Fig. 7). While their direct causal relationship remains unclear, subnuclear compartmentalization of the genome, i.e., physical separation of A and B compartments, occurs during early G1-phase as assayed by 4C-seq^19^ (<u>3C</u> <u>c</u>ombined with <u>seq</u>uencing), coincident with the establishment of a DNA replication timing program^48^. This result suggests that physical separation of A and B compartment sequences in early G1 might be involved in regulating replication timing in the upcoming S-phase. In contrast, precise radial subnuclear positioning is clearly not important for regulating replication timing, since homologous chromosomal domains exhibit nearly identical replication timing (Fig. 6a, b) and yet they often exhibit distinct radial subnuclear positions^27^. Rather, this remarkable similarity between homologous chromosomes may indicate that the additional regulatory information of DNA replication timing, and A/B compartments may in turn be largely coded in our genomic DNA.

This putative ‘genetically encoded’ regulatory mechanism, if it exists, may rely on physical interactions between different DNA sequences on the genome, starting with those between various *cis* regulatory elements within TADs. However, TADs and A/B compartments appear to be regulated independently, as factors required to maintain TAD organization are not required for A/B compartment separation^49,50^. Thus, this putative mechanism might guide TADs with similar properties to physically associate with each other to form A and B compartments, which in turn regulates replication timing but is not related to the formation of TADs per se. Interestingly, it has been shown that DNA fragments that underwent chromosomal rearrangements could retain their original DNA replication timing only if the fragments were more than 500-kb long^51^, suggesting that a certain threshold DNA length is necessary for this putative mechanism to exert its effect.

Such a mechanism may function at the level of binding of pre-replication complex (pre-RC) components to replication origins^46,52,53,54^ or replication origin density differences between early- and late S-phase compartments^55^. These properties of replication origins and replication timing may be ultimately governed by local chromatin ‘accessibility,’ for instance DNaseI hypersensitivity, which has been shown to serve as an excellent predictor of genome-wide DNA replication timing profiles by defining origin location and density^56^, although their causal relationship remains unclear.

Stochastic origin firing still occurs in the context of individual replication domains in mammalian cells^8,9,10^ and possibly within a given subnuclear compartment. However, sufficient density of potential origins in each domain, along with the additional regulatory layer discussed above, probably ensures nearly conserved replication timing between cells, cell cycles, and homologs (Figs. 3a, 6a–c), especially in early S-phase when the origin density is highest^52,55^. Because late-replicating domains show lower density of potential origins than early-replicating domains^52,55^, it would be interesting to investigate whether replication timing becomes more heterogeneous among cells toward the end of S-phase.

Transcription has been shown to correlate with replication timing^4,26^ and influence origin activity^57,58^. While single-cell RNA-seq studies have demonstrated stochasticity of gene expression in individual cells^59,60,61,62^, our results suggest that transcriptional heterogeneity between cells does not influence DNA replication timing much, which is consistent with the presence of an additional regulatory layer of DNA replication timing on top of stochastic origin firing. However, it is possible that subtle differences in replication timing exist due to gene expression stochasticity, but they may be difficult to detect in genomic regions of high density of potential origins and transcription units, i.e., early-replicating regions. In support of this idea, we noticed a subtle but detectable difference between homologs at an imprinted region at the *Grb10* locus, which is replicated around mid S, with the more active homolog replicating earlier than the less active one (Supplementary Fig. 4). Our observation is that imprinting regions are relatively gene-poor, and such regions might reveal the effect of transcription on origin firing more so than gene-rich, early-replicating regions with higher origin density. Thus, by sorting cells at different S-phase time windows, it might be possible to detect replication timing asynchrony of other imprinting regions or even discover novel imprinted regions with replication timing asynchrony.

As exemplified by allelic differences in the *Grb10* imprinting region and the X chromosomes in differentiated mESCs, the DNA replication timing program is largely but not entirely determined by DNA sequence. Moreover, we successfully detected differentiation-induced replication timing changes at the single-cell level (Fig. 2) and further discovered that these regions exhibit higher cell-to-cell heterogeneity than constitutive regions in mESCs (Fig. 4a). This finding suggests an intriguing possibility that sequences subject to developmental regulation have inherent instability of replication timing regulation under certain circumstances. For instance, these domains could be less well defined structurally by DNA sequence, unlike constitutively early- or late-replicating domains, and their inherent instability may confer competence for developmental regulation. Such inherent instability is necessary but not sufficient for developmental regulation; the actual changes must be induced epigenetically, such as in the case of XCI, where both X chromosomes may be competent but only one becomes inactivated and late-replicating (Fig. 5c). From a larger perspective, it is tempting to speculate that there is a link between acquisition of sequences prone to cell-to-cell heterogeneity and the emergence of multi-cellularity during evolution.

Cell-to-cell heterogeneity in replication timing of developmentally regulated D-class sequences, though only in mESCs, is reminiscent of the property of DNA sequences associated with the nuclear lamina, or LADs, which correlates well with late replication^63^. That is, fLADs, which show developmental regulation of nuclear lamina association, exhibit higher cell-to-cell heterogeneity than cLADs^64^. However, the degree of fLAD heterogeneity is much higher than that of the developmentally regulated D-class replication domains (Fig. 4), suggesting that they are probably distinct. Interestingly, D-class sequences show poorer subnuclear compartmentalization than constitutive classes during G1 when chromosomes reacquire the interphase structure^19^. This may be more pronounced in mESCs that show higher replication timing heterogeneity than differentiated cells and may be linked to the more ‘open’ chromatin structure in mESCs than in differentiated cells^65,66,67^.

In addition to significantly improving our current model of DNA replication and providing insights into 3D genome organization, our scRepli-seq technology is a valuable addition to the existing single-cell epigenome profiling methods, including BS-seq^68,69^, ATAC-seq^70,71^, DNaseI-seq^72^, ChIP-seq^73^, LaminB1 DamID^64^, Hi-C^40,41,42^ and more^74^. While these technologies opened doors to a single-cell era in molecular biology, many of them are still technically challenging. Because DNA replication domains are regulated at the Mb-scale, scRepli-seq does not require high read depth per sample (Fig. 2b). As a result, our scRepli-seq methodology generates affordable (<100 US dollars per sample) yet highly informative data sets that have sufficient resolution to visualize replication domains genome-wide in a manner comparable to cell population studies. Furthermore, the strong correlation between DNA replication timing and A/B compartments makes scRepli-seq a valuable alternative for interrogating the 3D organization of chromosomes in single cells, along with the single-cell Lamin B1 DamID technology^64^. In fact, our scRepli-seq results suggest an intriguing possibility that A/B compartment organization is also conserved from cell to cell and between homologous chromosomes.

Our scRepli-seq technology is versatile. In addition to its predictive power of A/B compartment organization, we also demonstrated the feasibility of concurrent CNA analysis to address chromosomal abnormalities (Table 1, Supplementary Fig. 5). We also believe that concurrent single-cell RNA-seq analysis is a very likely future option^75,76^ to assess the effect of transcription on replication at the single-cell level. Various imaging techniques, such as immunofluorescence and replication foci detection, should also be feasible prior to scRepli-seq. In addition, our method allowed identification of the Xi in a straightforward manner (Fig. 5), making it one of the most reliable techniques for obtaining haplotype-resolved genome-wide data (Fig. 6). We believe the remarkable simplicity of our scRepli-seq method makes it well suited for combining with other technologies in the future to gain novel insights into the regulation of DNA replication and the 3D genome organization at an unprecedented resolution for a single-cell methodology.

## METHODS

### Cell culture and mESC differentiation

hTERT-RPE1 cells were grown in MEM-alpha supplemented with 10% FBS and penicillin/streptomycin. CBMS1 mESCs have been described^20^ and were grown in 2i/LIF medium as described^77^. For differentiation, CBMS1 mESCs were differentiated to EpiLCs for 2 days and then switched to aggregation culture (EB/embryoid body culture) in Nunclon Sphera 96U-well plates (ThermoFisher, #174925), starting from 2,000 EpiLCs per well exactly as described^77^, except for the use of plain GK15 medium^77^ without any additional factors added during the aggregation culture. This process is practically identical to the SFEBq neural method of mESC differentiation (serum-free floating culture of EB-like aggregates with quick reaggregation)^78^, except that we started from EpiLCs instead of mESCs. In our hands, this resulted in efficient formation of neurectoderm cells based on gene expression after 7 days of differentiation (2 days to EpiLCs and then 5 additional days of EB culture). For FACS experiments, cells were fixed in 75% ethanol as described^5^ after singlecell suspension with trypsin for day-7 EBs^79^.

### Sample preparation for replication timing profiling of cell populations

We followed our routine BrdU-IP-based protocol as described^11^. For FACS, we used a Sony SH800 cell sorter in the ultra-purity mode, fractionating early and late S-phase populations. The BrdU-IP protocol has been described in detail^11^, except that we used a Bioruptor UCD-250 (Sonic Bio) for gDNA sonication in high output mode, with ON/OFF pulse times of 30 s/30 s for 6 min. After BrdU-IP, immunoprecipitated DNA samples were subject to WGA with a GenomePlex kit (Sigma, WGA2) for CGH microarray analysis^11^ and with a SeqPlex kit (Sigma, SEQXE) for NGS analysis. For CGH microarrays, we used the SurePrint G3 Mouse CGH 4x180K Array from Agilent (G4839A), labeling early and late-replicating DNA samples after WGA with Cy3 and Cy5 or vice versa followed by overnight hybridization, washing and slide scanning, according to the manufacturer’s instructions. For NGS analysis, NGS libraries were constructed from early and late-replicating DNA after WGA with an NGS LTP Library Preparation Kit (KAPA, KK8232) according to the manufacturer’s instructions and were subject to NGS with an Illumina Hiseq 1500 system (CBMS1 mESCs). For NGS analysis on an Ion Proton system (hTERT-RPE1), NGS libraries were constructed using Ion Plus Core Module for AB Library Builder System (Thermo Fisher Scientific, #4477683) with Ion Xpress Barcode Adapters 1-16 Kit (Thermo Fisher Scientific, #4477683) according to the manufacturer’s instructions. For a copy number-based analysis of the early S-phase population (Supplementary Fig. 1b), 200,000 cells from the first half of S-phase were sorted by FACS, and gDNA was isolated using a Qiagen kit (Qiagen #69504, DNeasy Blood & Tissue Kit) and fragmented to 200–300 bp with a Covaris ultrasonciator (model: S220, tube: microTUBE snap-cap) according to the manufacturer’s instructions (peak incident power: 175, duty factor: 10%, cycles per burst: 200, treatment time: 120), followed by cleanup and size selection via Agencourt AMPure XP SPRI bead treatment (Beckman Coulter; first and second round size selection, 0.6x and 1.8x reaction volume, respectively). The samples were then subjected to NGS library preparation and NGS as described above.

### Sample preparation for replication timing profiling of single cells and 100 cells

Single or 100 mid-S or G1 cells were sorted with a Sony SH800 cell sorter using the singlecell mode. Sample preparations were based on Baslan et al^16^. Single or 100 cells were sorted directly into a 96-well plate with 6 μl of cell lysis buffer [352 μl H_2_O, 1 μl of 10 mg/ml Proteinase K (Sigma, P4850), 16 μl 10x single-cell lysis and fragmentation buffer (Sigma, L1043)], incubated at 55°C for 1 h and then at 99°C for 4 min for gDNA isolation and fragmentation. Then, 6 μl of the gDNA solution was subjected to WGA with a SeqPlex kit (Sigma, SEQXE) in a 30-μl reaction volume as per the manufacturer’s instructions. Amplified gDNA was purified and size-selected with Agencourt AMPure XP SPRI beads (1.7x reaction volume), and the SEQXE adapter sequence was removed by the primer removal enzyme Eco57I (Sigma, SEQXE). The gDNA was then purified with Agencourt AMPure XP SPRI beads (2.0x reaction volume) and eluted in 20 μl of 1/10x Elution Buffer (Qiagen). The DNA fragment size peak should be within 150–200 bp, which was confirmed by a capillary electrophoresis system, MultiNA (Shimadzu). We could easily distinguish single cells from 0 or 2 cells by quantification of DNA by MultiNA, which provided reassurance that gDNA was indeed derived from single cells. Then, NGS libraries were constructed with an NGS LTP Library Preparation Kit (KAPA, KK8232) according to the manufacturer’s instructions, with slight modifications based on Kadota et al^80^. For a multi-plex NGS run, a SeqCap adapter kit A/B (Roche, 07141530001/ 07141548001) and NEXTflex DNA barcode (Bio Scientific, NOVA) were used. Finally, the samples were subjected to NGS on an Illumina Hiseq 1500 system (80-bp length, single-read or paired-end read).

### NGS read mapping and allele-specific mapping

The raw Fastq are files were trimmed to remove adapter sequences using the *cutadapt* program^81^ before mapping. For single-cell and 100-cell Repli-seq, we performed a two-step adapter trimming, first removing the Illumina adapter based on the index of each NGS library and then removing the SEQXE adapter. As the SEQXE adapter sequence was not available, we empirically estimated it as the sequence that repeatedly appeared near the 5’ end. Mouse and human reference genomes, mm9 [chr1–19, chrX, chrM, and chrR (one copy of rDNA, GenBank: BK000964.1)] and hg19 (chr1–22, chrX, chrM) assemblies, were used. For haplotype-resolved analysis, we constructed the CBMS1 mESC-specific diploid genome as described in Sakata et al.^82^ with minor modifications: (i) fermi v1.1-r751 (Li, 2012) with default options was used for *de novo* assembly of MsM genomic reads; (ii) the maximum indel length was 30 bp; (iii) only variants located at informative positions between the CBA and MsM strains were considered. The NCBI Sequence Read Archive accession numbers of strain-specific genomic reads used were DRP000194 (MsM) and ERP000927 (CBA; only library 3888059)^21,83^. For mapping, *bwa*^84^ (ver: 0.7.10-r789) was used (command: bwa aln => bwa samse). For mapping to mm9 or hg19 reference genomes, we used the picard tool (http://broadinstitute.github.io/picard/) to remove duplicated reads and defined MAPQ>10 as uniquely mapped reads. For mapping to the CBA/MsM diploid genome, we defined MAPQ>16 as allele-specific reads and used the liftover tool (UCSC Genome Browser) to convert to the mm9 genome coordinates. Among the reads converted to mm9 coordinates, we filtered out duplicated reads that had an identical chromosome start position and strand information relating to an existing read. We also filtered out reads that overlapped with the hg19 and mm9 black lists^85^ (https://sites.google.com/site/anshulkundaje/projects/blacklists).

### Computations associated with the replication timing profiling of cell populations

After mapping, we followed an established standard analytical procedure for BrdU-IP population replication timing (RT) analysis using CGH microarrays^11^. For BrdU-IP population RT analysis by NGS, we counted the reads of early and late S-phase BrdU-IP samples in sliding windows of 200 kb at 40-kb or 80-kb intervals or in non-overlapping 400kb windows and performed rpm (reads per million) normalization. Then, the ratio of early-S to total read counts [(Early-S reads)/(Early-S reads + Late-S reads)] was calculated for each bin and their distribution converted to fit within a ± 1 scale, and this value was defined as the BrdU-IP RT score of each bin. We filtered out bins whose total read counts were within the bottom 5% of all bins. For haplotype-resolved RT profiling, we followed the exact same procedures using a 400-kb bin size. To convert the BrdU-IP RT values to the ‘% S-phase’ values used in Figs. 3, 4, and 6, we ranked the BrdU-IP RT values of all 80-kb bins (sliding windows of 200 kb at 80-kb intervals) or 400-kb bins throughout the genome from the earliest to the latest, and assigned the percentile rank of each bin as its % S-phase value, and in these figures, we subdivided the genomic bins into one-percentile groups based on their % S-phase value. For analysis of 200,000 early S-phase and G1 control cell populations (Supplementary Fig. 1b), we counted the reads of early-S and G1 in sliding windows of 200 kb at 40-kb intervals, performed rpm normalization in a manner identical to BrdU-IP NGS data processing, and defined Log_2_[(Early-S reads)/(G1 reads)] as the population early-S RT score. For analysis of NGS data derived from 100 cells, we counted the reads of 100 mid-S and G1 cells in sliding windows of 200 kb at 40-kb intervals and used the *correctMappability* command in the R package AneuFinder^86^ (http://bioconductor.org/packages/release/bioc/html/AneuFinder.html) for normalizing mid-S data based on G1 data. From the mappability corrected mid-S read counts, the genome-wide median was obtained and used to generate Log_2_[(Mappability corrected Mid-S reads)/median] scores, which we defined as the 100-cell mid-S RT score. Tag density values are defined as the read count per window divided by the total read count.

### Computations associated with the replication timing profiling of single cells

For quality control of scRepli-seq data, we reasoned that using median-absolute-deviation (MAD) scores is a simple and effective approach to filter out cells with problematic RT data distribution because G1 and mid-S cells are expected to show relatively small and large RT variability, respectively. For each cell, a MAD score was calculated for Log2[(counted reads)/(genome-wide median of counted reads)] in non-overlapping 200-kb windows, and we empirically filtered out cells with MAD scores of >0.3 for G1 cells and of <0.4 and >0.8 for mid-S cells. The number (ratio) of cells in each group that passed this criteria was as follows: 4/4 (G1, hTERT-RPE1), 14/20 (mid-S, hTERT-RPE1), 5/5 (G1, mESCs), 34/36 (mid-S, mESCs), 6/7 (G1, day 7), and 43/52 (mid-S, day 7). For the G1 cells, we merged multiple (i.e., 3–5) cell samples and used these data as control data. To select high-quality G1 cells, we discarded samples with chromosomal instability, which was possible by using the *findCNVs* command in AneuFinder^86^ with a 500-kb bin size [6-HMM options: method-?MM”, max.iter=3000, states=c(“zero-inflation”, “0-somy”, “1-somy”, “2-somy”, “3-somy”, “4-somy”, “5-somy”, “6-somy”), eps=0.01]. As a result, 3/4 hTERT-RPE1 cells, 5/5 CBMS1 mESCs, and 3/6 day-7 cells in G1-phase had identical karyotypes and were merged to generate a control G1 single-cell data set in each cell type. For the analysis of single mid-S cells, we counted the reads in sliding windows of 200 kb at 40-kb intervals, and used the AneuFinder’s *correctMappability* command^86^ for normalizing mid-S data based on the merged G1 control. From the mappability corrected mid-S read counts, the genome-wide median was obtained and was used to generate Log_2_[(Mappability corrected Mid-S reads)/median] scores, which we defined as the single-cell mid-S RT score. At this point, one day-7 cell showed an uninterpretable profile upon visual inspection on the IGV browser and was discarded from further analysis. For haplotype-resolved RT analysis of CBMS1, we applied the AneuFinder’s*findCNVs* command^86^ to CBA and MsM data at 1-Mb bins [6-HMM options: method=“HMM”, max.iter=3000, states=c(“zero-inflation”, “0-somy”, “1-somy”, “2-somy”, “3-somy”, “4-somy”), eps=0.01]. We did not find major chromosomal instabilities in mESCs (except for trisomy 8, which will be discussed below), but variable results were seen on chromosome 18 in day-7 G1 cells, with two cells having both chromosome 18 derived from CBA while one had both CBA- and MsM-derived chr18 as expected. As a result, we excluded chromosome 18 from haplotype-resolved analysis. We immediately noticed that haplotype-resolved, CBA- and MsM-specific data sets exhibited higher variability of read counts across the genome than non-resolved mm9 data. For instance, X-chromosome read coverage was lower than autosomes, probably due to the lower SNP density than that of autosomes, and some autosomes had sequences that show higher read coverage than others, likely due to duplication. We therefore subdivided the genome into 3 groups based on these read count differences in the merged G1 control: (1) normal coverage bins on autosomes [(CBA in mESCs: chr1–7, 9-19; MsM in mESCs: chr1–19; CBA in day-7 cells: chr1–11, 12 (a single copy region), 13-17, 19; MsM in day-7 cells: chr1–17, 19), (2) high coverage bins on autosomes [CBA in mESCs: chr8; CBA in day-7 cells: chr12: 75,000,001-121,000,000 (a duplicated portion of chr12)], and (3) chrX. Then, we calculated the median ± 1.5x IQR (interquartile range) for each group and filtered out highly variable bins that were outside this range (filtered and merged G1 control). After these procedures, we used the AneuFinder’s *correctMappability* command^86^ for normalizing mid-S data based on the filtered and merged G1 control in sliding windows of 1 Mb at 40-kb intervals and generated Log_2_[(Mappability corrected Mid-S reads)/median] scores, which we defined as the haplotype-resolved single-cell mid-S RT score. Regarding trisomy 8, we found that 97% (33/34) of our mid-S CBMS1 mESCs exhibited trisomy 8. In contrast, our day-7 cells were largely non-trisomy 8 (3/42=7.1%) because we used CBMS1 mESCs passaged ~1 month earlier for differentiation. This result suggests that trisomy 8 can quickly overwhelm the mESC population, while it does not occur rapidly during mESC differentiation.

### Pearson correlation matrix, hierarchical clustering and tSNE analysis of single-cell Repli-seq data

Single-cell mid-S RT scores (in sliding windows of 200 kb at 40-kb intervals) within the median ± 1.5x IQR range, excluding the X-chromosome bins, were used to generate a Pearson correlation matrix and for hierarchical clustering analysis using Ward’s method. For tSNE analysis, we used the R package RtSNE (https://github.com/jkrijthe/Rtsne) with the default setting using sliding windows of 200 kb (or 1 Mb for haplotype-resolved assay) at 40kb intervals. For the analysis of the X chromosomes in Fig. 6, we limited the analysis to cells that retained two X chromosomes (31 mESCs and 26 day-7 cells).

### Binarization of scRepli-seq data

Binarization was performed using the Mappability corrected Mid-S reads described above by using the *findCNVs* command in AneuFinder^86^. For haplotype-unresolved analysis, nonoverlapping 80-kb/400-kb windows were analyzed [2-HMM; options: method=“HMM”, max.iter=3000, states=c(“zero-inflation”, “0-somy”, “1-somy”, “2-somy”), eps=0.01; 1-somy, unreplicated; 2-somy, replicated]. For haplotype-resolved analysis, non-overlapping 400-kb windows were analyzed [2-HMM; options: method=“HMM”, max.iter=5000, states=c(“zero-inflation”, “0-somy”, “1-somy”, “2-somy”), eps=0.01].

### Computation of variability scores and average replication timing scores of single cells

Cell-to-cell variability scores, ranging from 0 to 1, were calculated for each genomic bin by comparing binarized scRepli-seq data across cells. The variability score of a given bin was 0 when it was either replicated in all cells or unreplicated in all cells, while the score was 1 when it was replicated in 50% of all cells analyzed. “N/A” cells were excluded from the analysis. To calculate within-cell variability scores, genomic bins were first subdivided into one-percentile groups based on their BrdU-IP RT scores, with the earliest- and latest-replicating bins corresponding to 0–1% and 99–100% S-phase groups, respectively. Within each group, we calculated the rate of bins that were replicated vs. unreplicated and converted this value to fit within 0 to 1, nearly identical to the cell-to-cell variability calculation. To calculate single-cell average RT scores, the rate of replication of a given genomic bin among the cell population analyzed was calculated, from which the mean RT of a given cell was subtracted for normalization.

### Fitting a Gaussian model to cell-to-cell variability score distribution

For both cell-to-cell and within-cell variability distribution, we fit a Gaussian model based on Equation 1 (Eq1), where a, b, and c are scaling factor, cell-to-cell/within-cell variability score, and SD, respectively. Start conditions were: a=1, b=50, and c=10 [nls(y~(a*exp(-0.5*(x-b)^Λ^2/(c^Λ^2))), start=list(a=1,b=50, c=10))]. We calculated the mean (b) and SD (c) values using the *nls* (nonlinear least squares) command of an R package, stat (x, % S-phase score; y, cell-to-cell or within-cell variability).

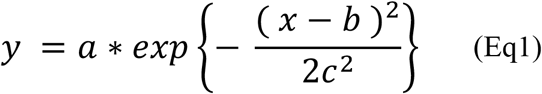

### Classification of mouse genomes into constitutively early (CE), constitutively late (CL), and developmentally regulated (D) sequences

A total of 28 mouse BrdU-IP RT data sets were analyzed at an 80-kb window size based on the method described by Dileep et al^19^. Definitions of the CE-, CL-, and D-class bins were identical to the definitions of Dileep et al^19^, except that our analysis was at an 80-kb bin size (Supplementary Fig. 2). Bins with BrdU-IP RT scores of >0.5 in all 28 data sets and <–0.2 in all 28 data sets were defined as CE- and CL-class bins, respectively. Bins with the maximum and the minimum BrdU-IP RT scores of >0.5 and <–0.5 among the 28 data sets, respectively, were defined as D-class bins. For CE vs. D and CL vs. D comparisons, sequences with BrdU-IP population RT scores within 0–30% and 50–100% S-phase ranges, respectively, were analyzed in one-percentile groups of the BrdU-IP population RT (i.e., mean variability score of each bin), followed by a permutation test (10,000 times) to calculate the p-values. Definitions of the D-class subcategories used in Fig. 4e–g are briefly described in the figure legend. However, more specifically, among the D-class bins that are early-replicating in mESCs, D (EtoL) is a subset that undergoes EtoL RT changes upon 7 days of differentiation [i.e., RT scores change from 0–30% (E) to 50–100% S-phase (L)], while D (EtoE) is a subset excluding D (EtoL). Likewise, among the D-class bins that are late replicating in mESCs, D (LtoE) is a subset that undergoes LtoE RT changes upon 7 days of differentiation [i.e., RT scores change from 50–100% (L) to 0–30% S-phase (E)], while D (LtoL) is a subset excluding D (LtoE).

### Identification of XO cells from Mid-S scRepli-seq data

After binarization of haplotype-resolved scRepli-seq data from mid-S cells, the percentage of covered genomic bins on X chromosomes derived from CBA and MsM strains were calculated in each cell. The cells with one of the two X chromosomes having a percentage of covered genomic bins <1% were defined as XO cells.

### Identification of allelic replication timing difference between homologs

We performed one-way ANOVA to compare BrdU-IP RT data sets with a 400-kb bin size, and bins with p<0.05 were deemed significantly different. In the case of binarized scRepli-seq data comparison, we performed Fisher’s exact test, and bins with p<0.05 were deemed significantly different.

### Hi-C and A/B compartment calling

Hi-C data derived from J1 mESCs were used for the analysis^47^. Cool format Hi-C fragment data sets (https://github.com/mirnylab/cooler) were downloaded (ftp://cooler.csail.mit.edu/coolers, file name: Dixon2012-J1 mESC-HindIII-allreps-filtered.frag.cool) and this .cool format fragment file was converted to 400-kb bin resolution data sets using the *cooler* tools and an in-house script. Normalization was performed using the *balance* command of *cooler* using default parameters. After this, A/B compartments were calculated using the *cworld* package (https://github.com/dekkerlab/cworld-dekker;matrix2compartment.pl, option: --ez).

### Data availability

All replication timing profiles (BrdU-IP, 100 cells and single-cell Repli-seq) generated in this study are deposited in the NCBI Gene Expression Omnibus database (GEO; http://www.ncbi.nlm.nih.gov/geo/).

## ACKNOWLEDGMENTS

We thank S. Kuraku and members of his laboratory for assistance with NGS and A. Tanigawa for technical assistance. We also thank D.M. Gilbert for exchanging unpublished observations, H. Niwa and K. Araki for CBMS1 mESCs, and B.D. Pope for helpful discussions. The authors apologize to colleagues whose work could not be cited owing to length limitations. This work was supported by RIKEN CDB intramural grant to I.H., the Special Postdoctoral Researcher (SPDR) Program of RIKEN to S.T., and a Grant-in-Aid for Scientific Research on Innovative Areas (JP16H01405) from the Ministry of Education, Culture, Sports, Science, and Technology (MEXT) to S-i.T.

## AUTHOR CONTRIBUTIONS

S.T., H.M., T.S., S-i.T., and I.H. conceived the project. S.T., T.S., and S-i.T. developed and conducted scRepli-seq and BrdU-IP experiments. S.T. and I.H. performed mESC culture, differentiation and sample collection. T.S. and S-i.T. performed hTERT-RPE1 cell culture and sample collection. K.N. and C.O. constructed diploid reference genome and helped with the haplotype-resolved analysis pipeline setup. H.M. and S-i.T. performed bioinformatics analyses. K.O. and M.O. supported for the design and execution of the project. S.T., H.M, S-i. T., and I.H. wrote the manuscript.

## SUPPLEMENTARY FIGURE LEGENDS

**Supplementary Figure 1.**
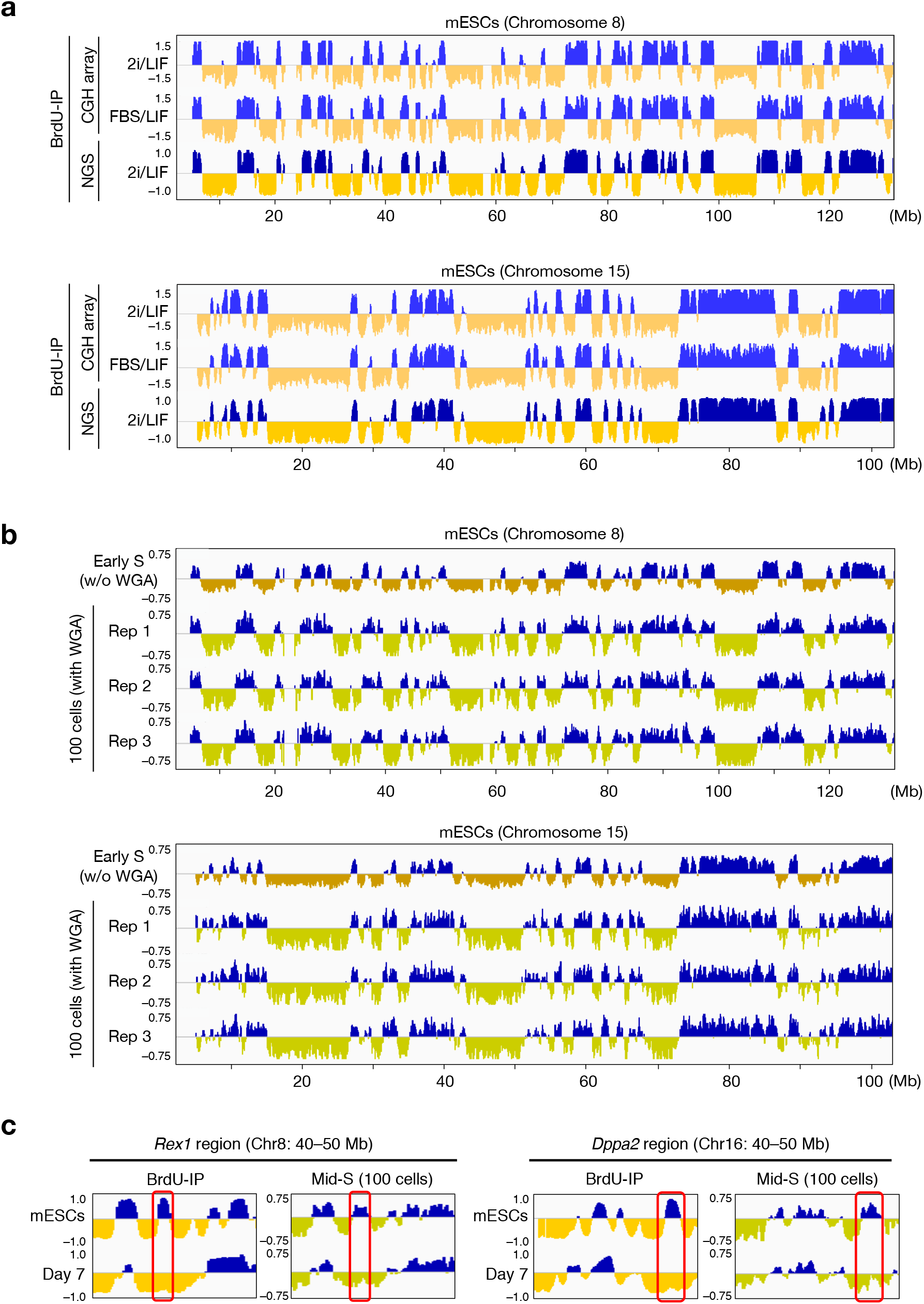
A copy number method generates replication timing profiles similar to those of previous methods. (**a**) BrdU-IP population RT analysis of mESCs cultured under 2i/LIF or FBS/LIF conditions, using CGH microarray or NGS, generated comparable profiles. Shown are mouse chromosomes 8 and 15. (**b**) A copy number-based RT analysis of early S-phase mESCs without WGA and three 100 mid-S mESCs with WGA generated comparable profiles. Shown are three replicates of 100 cells along with the early-S population on mouse chromosomes 8 and 15. (**c**) *Rex1* and *Dppa2* regions, which show EtoL changes upon mESC differentiation based on BrdU-IP population RT analysis, also showed the expected EtoL changes by copy number RT analysis of 100 mid-S cells.

**Supplementary Figure 2.**
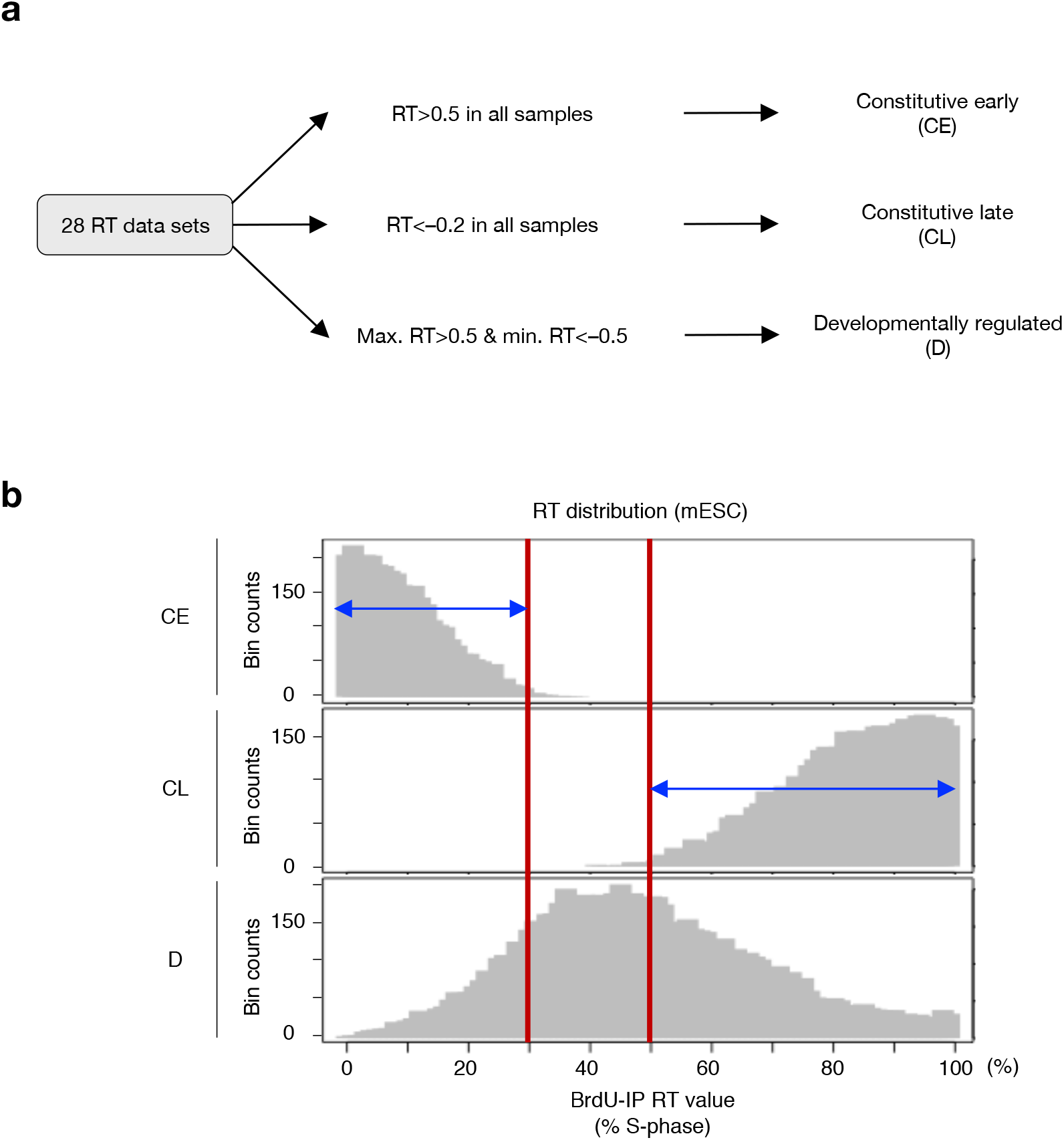
Definitions of constitutively early-replicating (CE), late-replicating (CL), and developmentally regulated (D) classes of sequences. (**a**) BrdU-IP population RT data from 28 types of cells were analyzed as shown in the flowchart to define CE-, CL- and D-class regions in a manner similar to that described by Dileep et al^19^. (**b**) RT distribution of each class based on BrdU-IP population RT data is shown. For CE vs. D and CL vs. D comparisons, sequences with BrdU-IP population RT values within 0–30% and 50–100% S-phase ranges, respectively, were analyzed in Fig. 4.

**Supplementary Figure 3.**
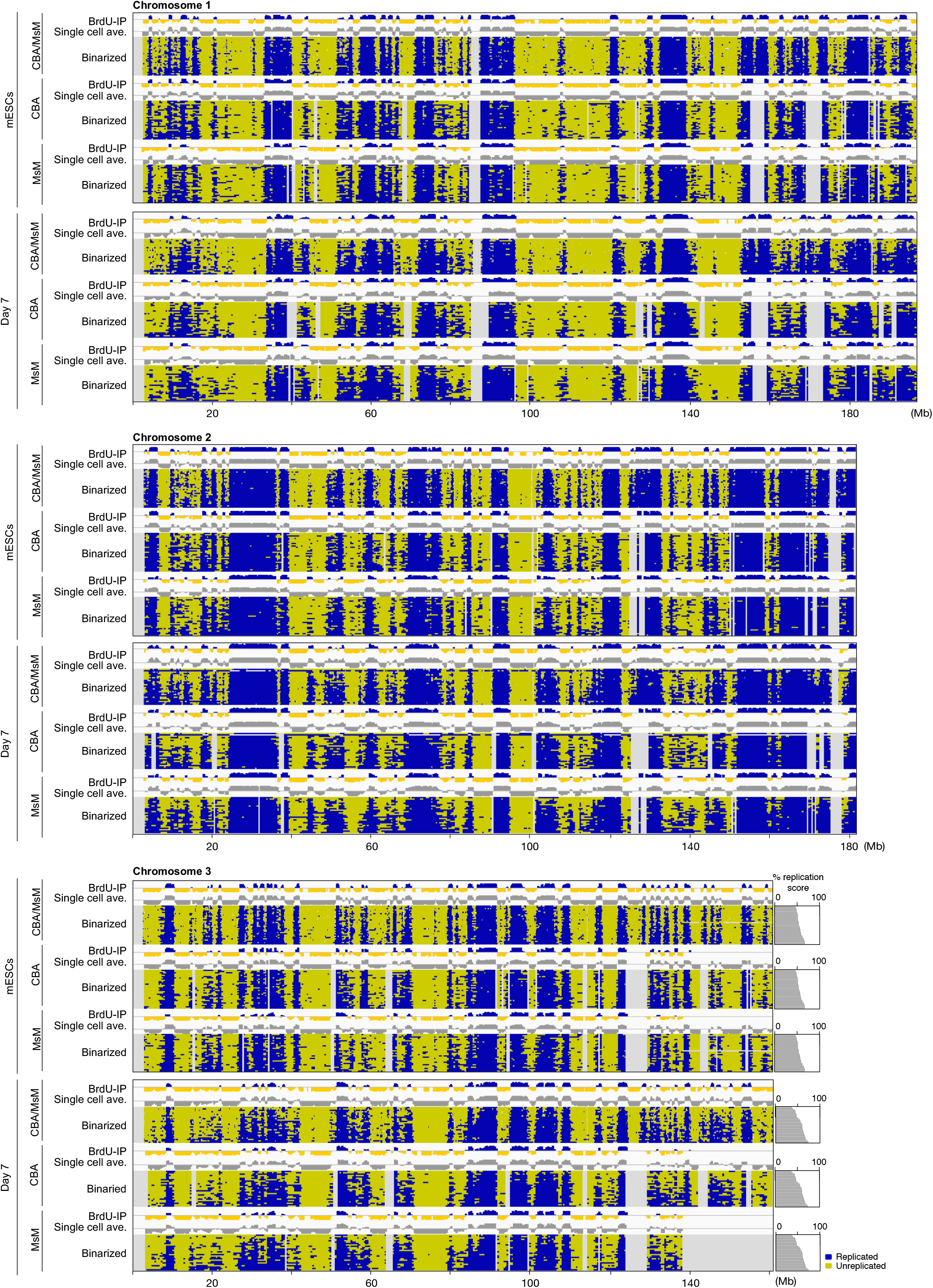
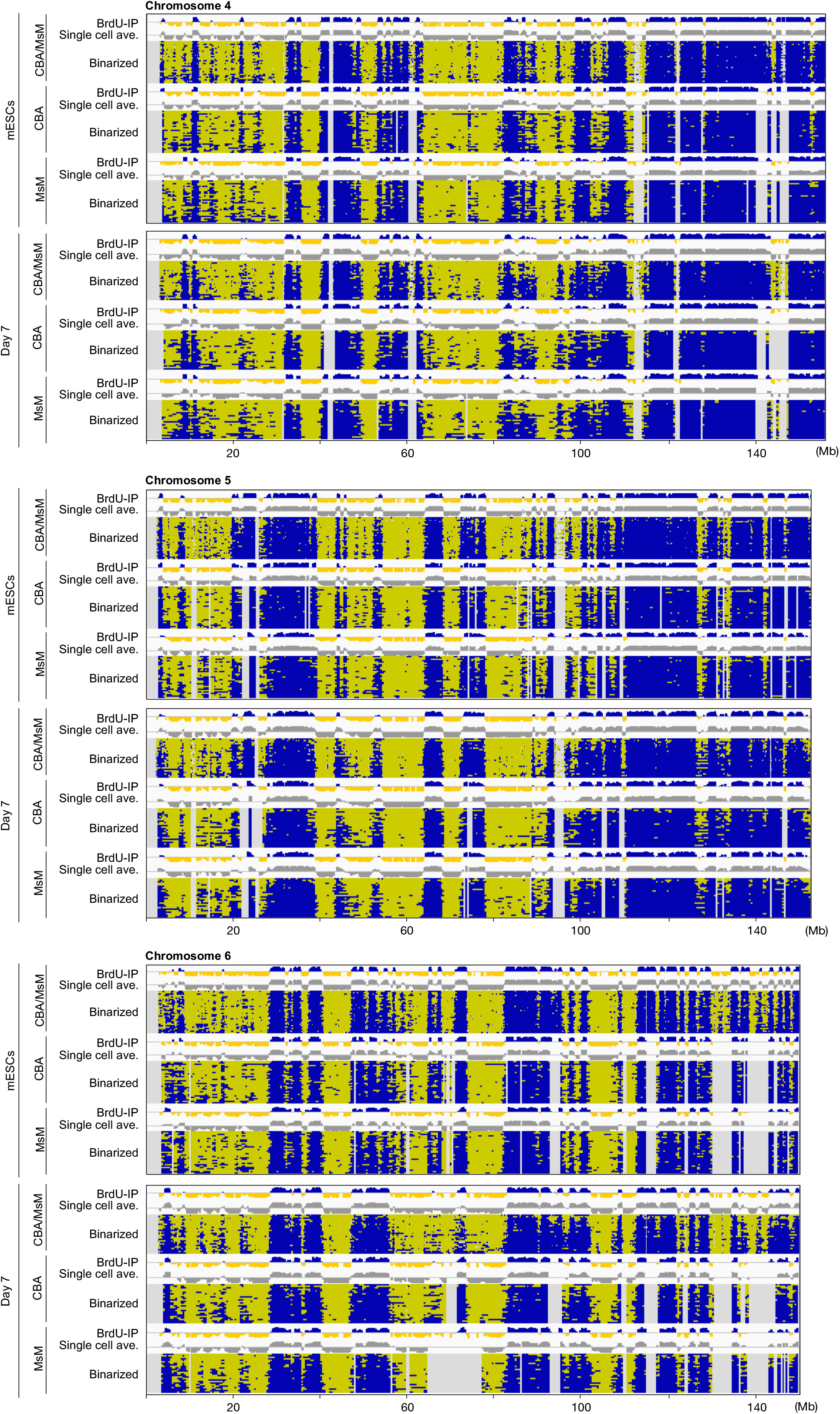
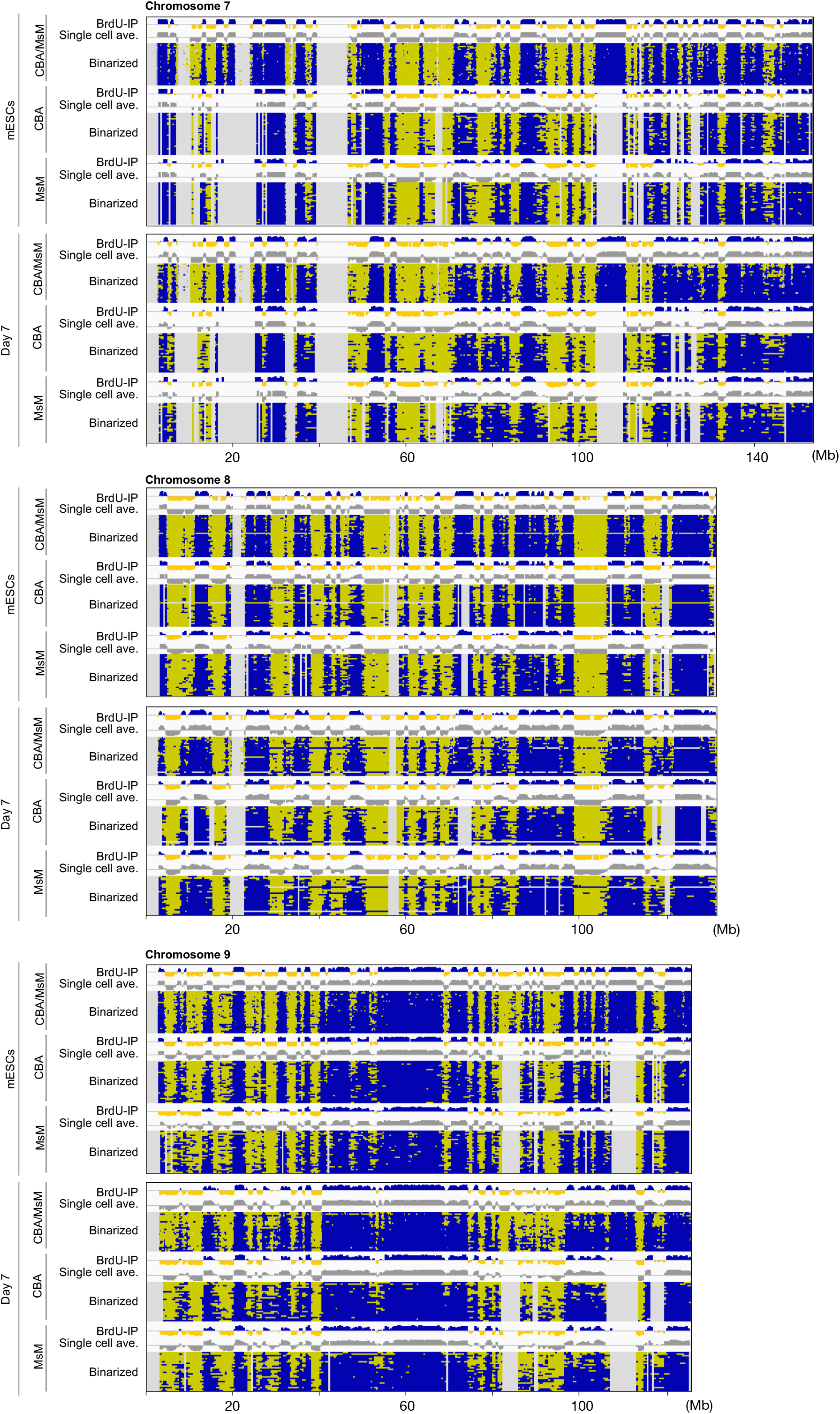
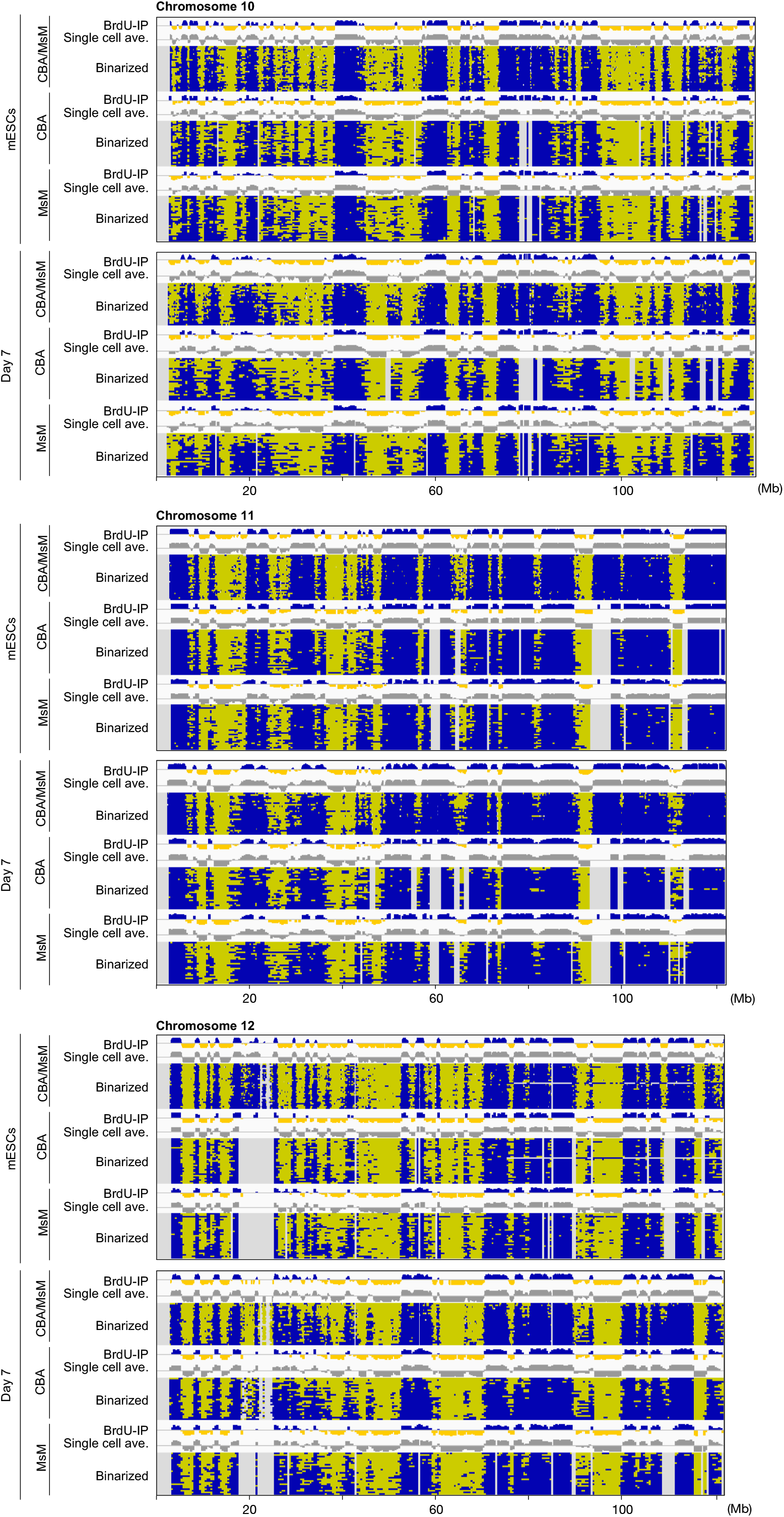
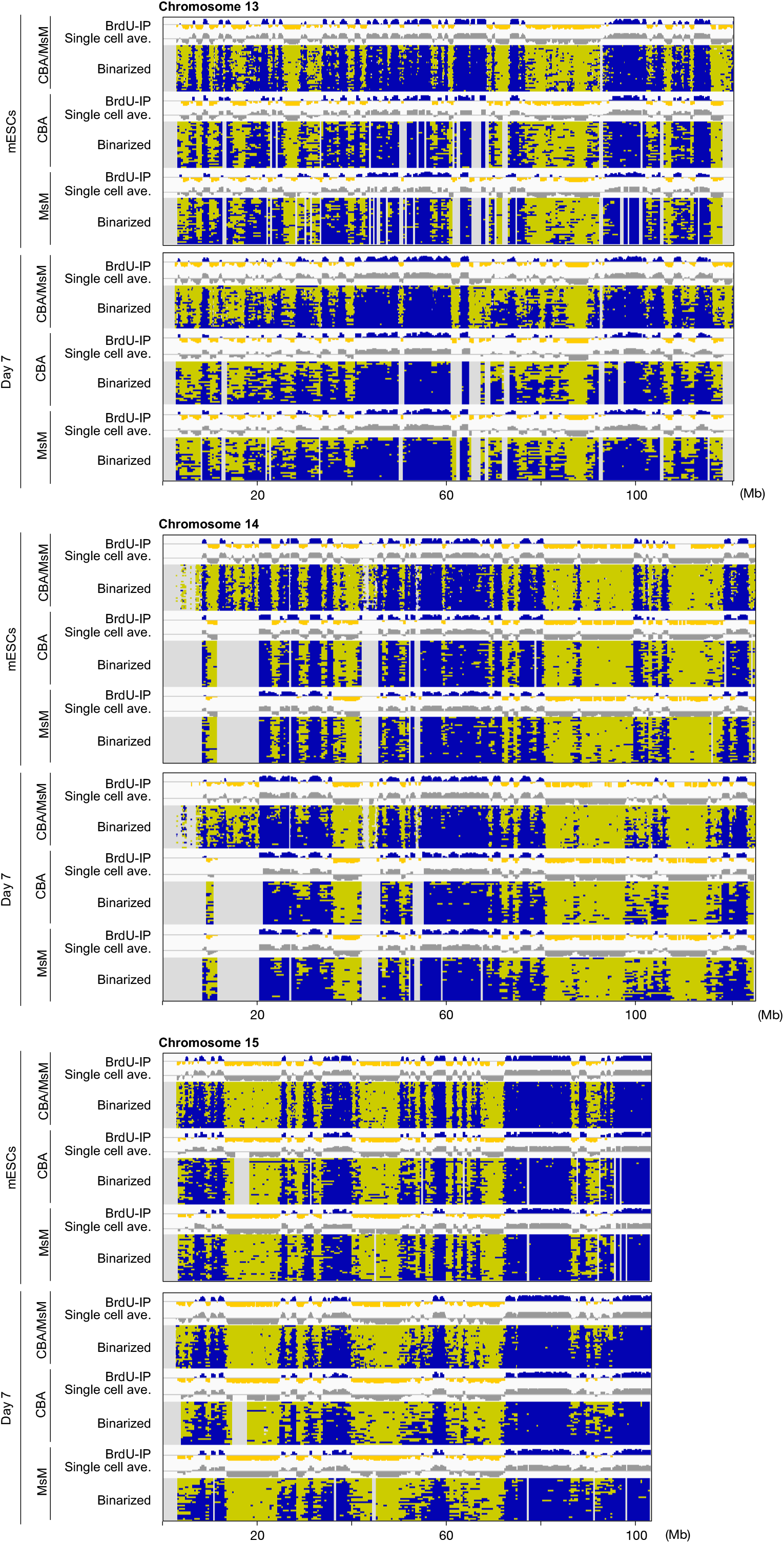
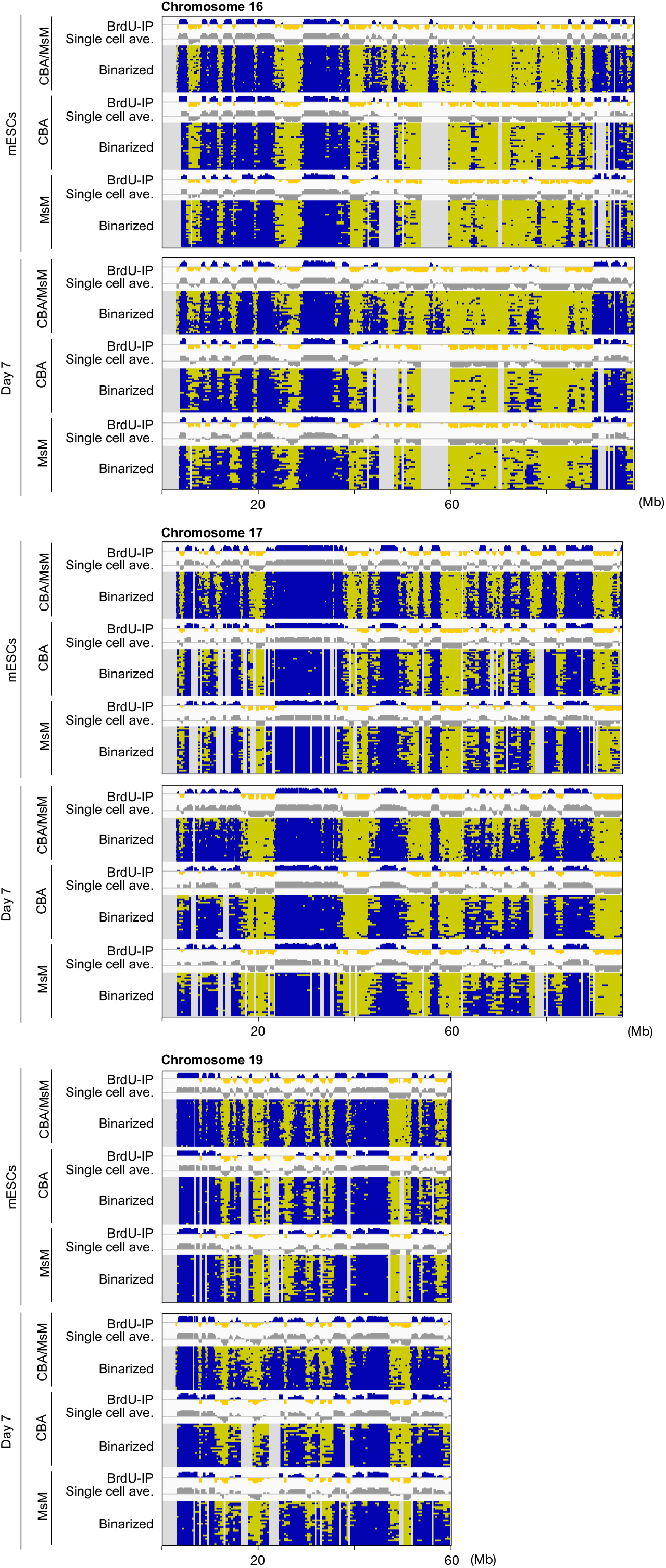
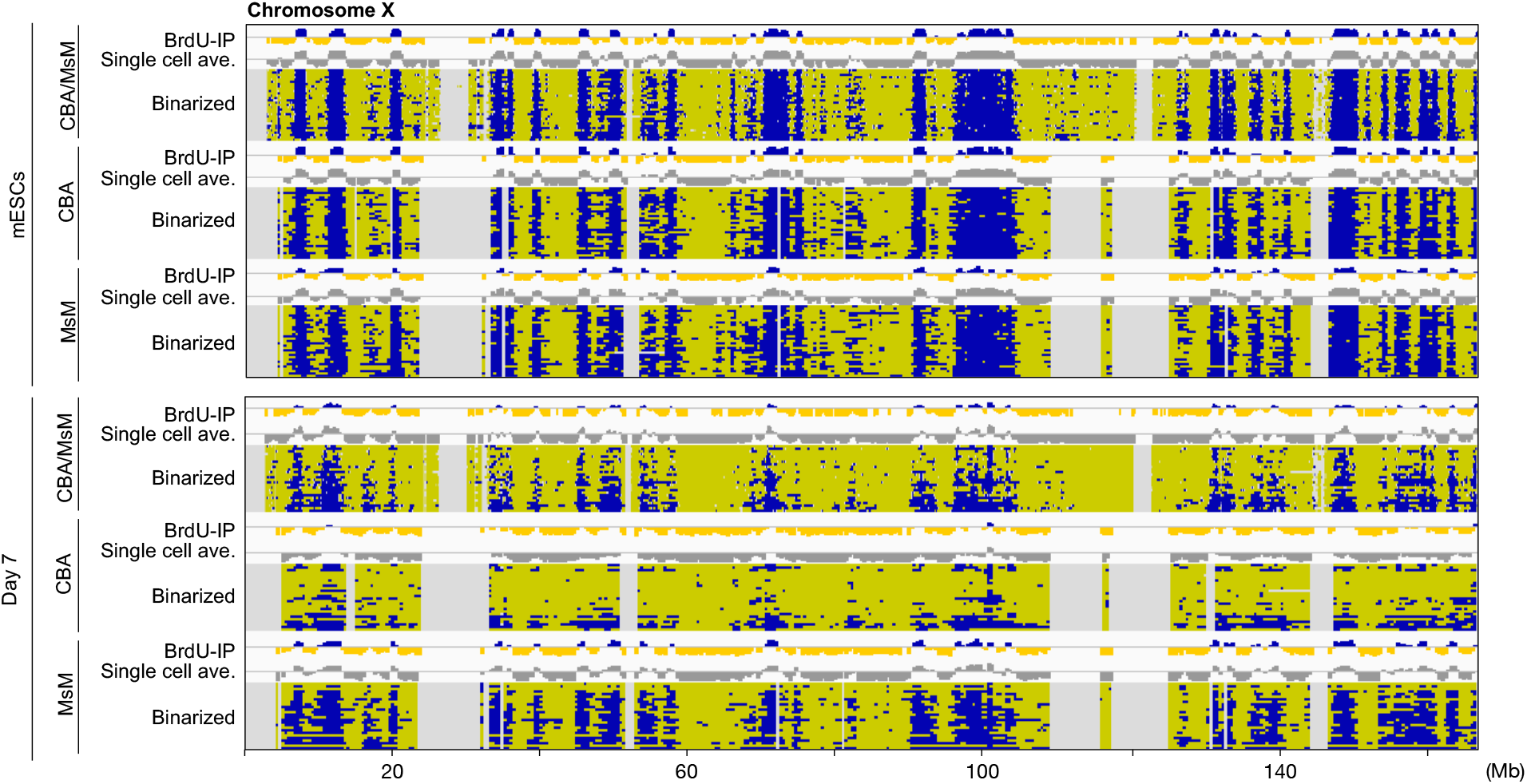
Binarized scRepli-seq profiles in mESCs and day-7 differentiated cells. Binarized scRepli-seq profiles with and without haplotype resolution on autosomes and the X chromosome, along with BrdU-IP population RT profiles and single-cell average profiles, are shown. Chromosome 18 is not shown, due to variable chromosomal losses and gains. Gray areas represent mapping failures. mESCs (31) and day-7 cells (26) are shown and are ordered according to their % replication scores, which are shown next to chromosome 3 profiles.

**Supplementary Figure 4.**
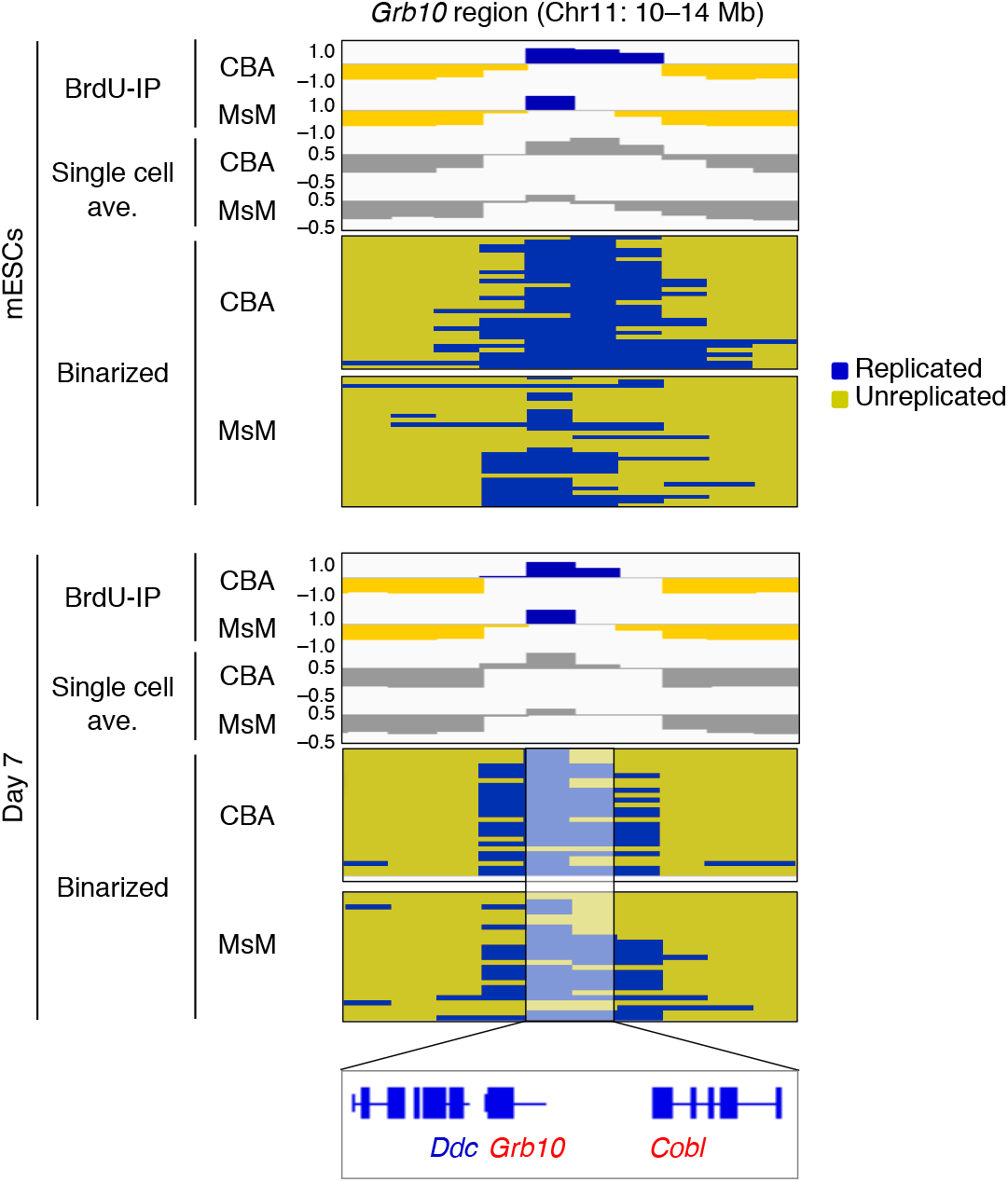
A magnified view of replication timing profiles surrounding the *Grb10* imprinting region on chromosome 11. Binarized and haplotype-resolved scRepli-seq profiles surrounding *Grb10,* along with BrdU-IP population RT profiles and averaged single-cell profiles, before and after mESC differentiation. Genes in blue and red are paternally and maternally expressed, respectively. Each pixel represents 400 kb.

**Supplementary Figure 5.**
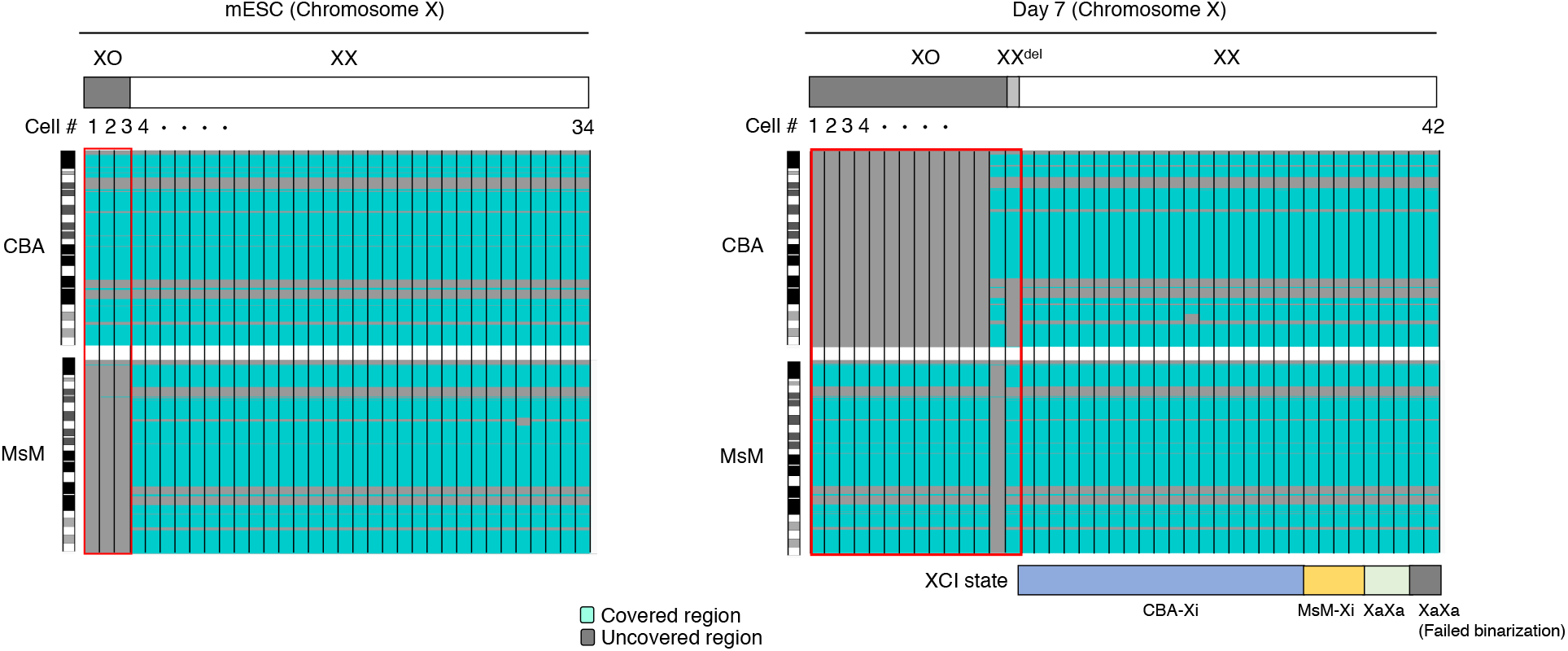
X-chromosome karyotypes in single cells. Schematic diagrams of the X-chromosome karyotype in 34 mESCs and 42 day-7 cells. Turquoise and gray represent covered and uncovered genomic regions, respectively. There were 13 XO, 1 XX^del^ (distal deletion of CBA-X), and 28 XX cells. XCI states of day-7 cells are also shown; ‘others’ represents samples that exhibited an Xa/Xa state based on tSNE analysis, and we categorized them as others due to the failure of binarization calling. ‘XaXa (failed binarization)’ represents samples that exhibited an Xa/Xa state based on tSNE analysis but showed binarization failures.

**Supplementary Figure 6.**
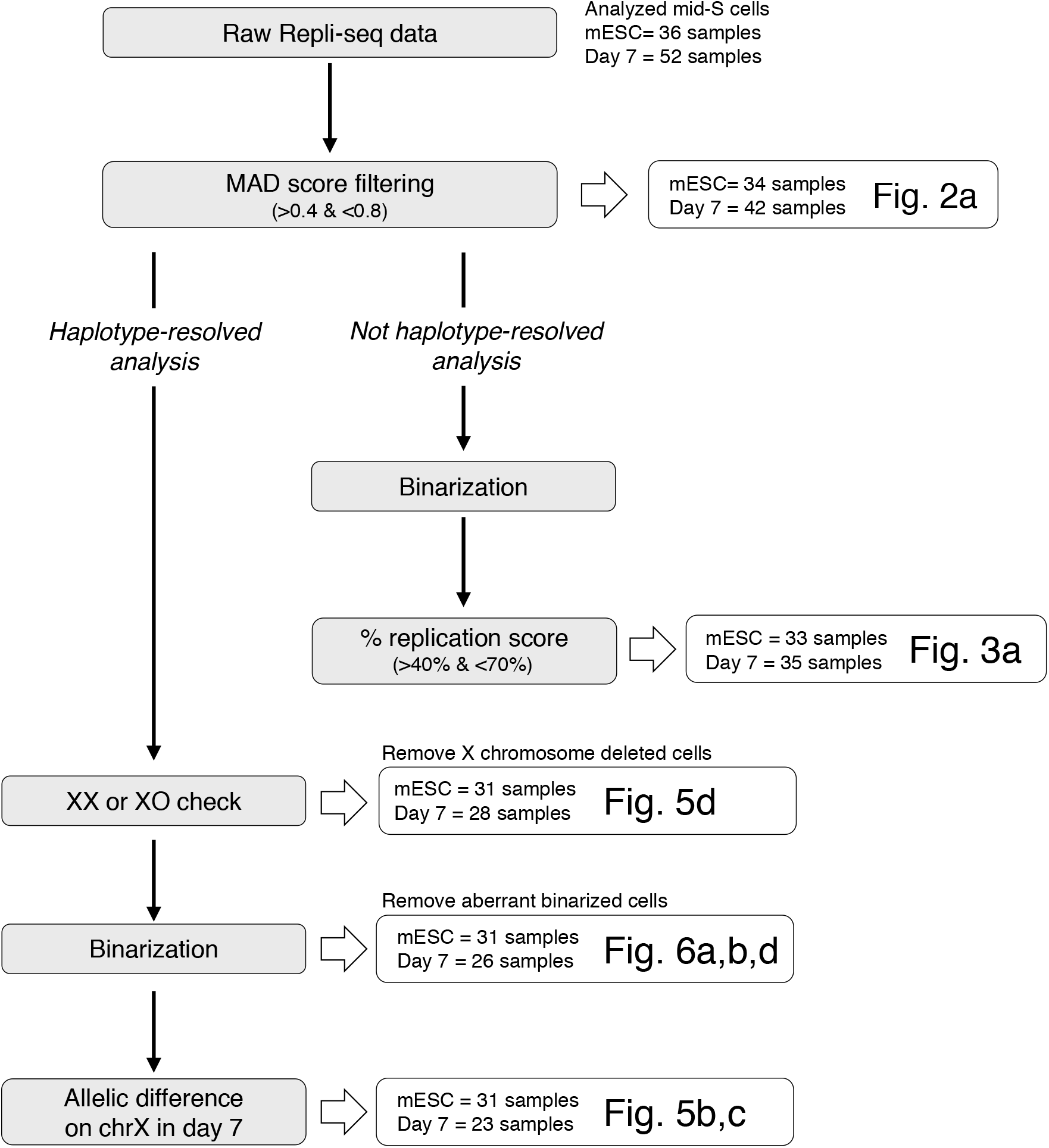
An overview of scRepli-seq analysis from raw data to various processed forms. A flow chart showing the fate of raw scRepli-seq data from 36 mESCs and 52 day-7 cells during various analytical procedures. For intermediate stage data, corresponding figure numbers are shown. Raw scRepli-seq data sets underwent MAD filtering to avoid data sets containing data variability below or above certain thresholds within a sample. For analysis without haplotype resolution, data sets that passed the MAD filtering were subject to binarization and those that passed the % replication range shown were analyzed. Otherwise, the data sets were haplotype-resolved, binarized and subjected to the same % replication filtering used for non-haplotype analysis. For the inactive X-chromosome analysis shown in Fig. 5, only 23 cells with a late-replicating X chromosome were analyzed.

